# An AI agent guides a pipeline for selective protein interaction inhibitors from discovery across targets to preclinical evaluation

**DOI:** 10.64898/2026.11.23.768516

**Authors:** Somaya A. Abdel-Rahman, Moustafa T. Gabr

## Abstract

A protein can engage multiple partners, making it challenging to disrupt disease-associated interactions while preserving others. We developed Bivexia, an AI agent that guides a multistage pipeline from discovery and de novo design through preclinical evaluation. Its decision rules are fixed in advance, and each prospective selection by the agent is cryptographically committed before testing. The objective is to discover small molecules that disrupt specified interactions of a target protein while preserving others. A compound advanced by Bivexia bound LAG-3 with nanomolar affinity and disrupted its interactions with tau and amyloid-β while preserving binding to MHC class II and FGL1. It reduced tau and amyloid-β burden in independent cohorts across two mouse models. The same rules gave three of nine hits on PrPᶜ and two of nine on LRP1, none in random controls. Our findings establish a framework for AI agents to pursue defined interaction selectivity across targets.

## INTRODUCTION

Drug selectivity can be defined at the level of the target protein or at the level of its individual interactions. For a protein with several binding partners, the goal may be to disrupt a specified subset of interactions while preserving others. Discovering drug candidates with this profile requires measuring both effects and using those measurements to decide which compounds advance. Computational models increasingly guide which molecules are synthesized and tested. A generative model proposed DDR1 inhibitors that were synthesized and validated within weeks (1), and deep learning identified structurally distinct antibiotics by screening libraries containing more than 10^8^ molecules (2, 3). AI has expanded access to structural models of drug targets (4), and compounds designed with machine learning assistance have entered clinical trials (5). AI agents extend this role by coordinating several steps of discovery, including generating hypotheses, proposing experiments, and interpreting results to guide follow-up studies. For example, the multi-agent system Robin combined literature analysis and experimental data interpretation to nominate and evaluate a drug repurposing candidate for dry age-related macular degeneration (6). Evaluating decisions made by an agent requires specifying in advance what a compound must achieve and which measurements will determine whether it advances. Retrospective benchmarks, random selection, and selections made by medicinal chemists address different questions about performance.

Retrospective performance can depend strongly on how the evaluation is conducted. A review of machine learning applications across 17 fields documented data leakage affecting 294 papers (7). In cheminformatics, random splits can place close analogs in both training and test sets, inflating apparent performance relative to temporal or scaffold splits (8, 9). Standardized benchmarks have helped make the novelty and validity of generated molecules comparable across studies (10, 11), although retrospective enrichment provides limited evidence of prospective performance (12). Feature attribution can explain which inputs influenced a prediction without establishing its experimental accuracy (13), and activity cliffs illustrate how small structural changes can produce large differences in potency (14).

A proposed framework for AI-assisted drug discovery treats model predictions as testable hypotheses and calls for documenting the rationale for testing each prediction, the expected result, and how alternative outcomes would guide the next decision (15). Work on evaluating drug discovery agents has similarly distinguished the assessment of task execution and output quality from testing whether an agent makes a scientific discovery (16). Clinical research provides a precedent for prospective evaluation through preregistration of outcomes, analyses, and stopping rules (17, 18). We apply this principle to compound advancement by fixing the decision rules before testing and cryptographically committing each prospective selection, so that subsequent changes are detectable. Releasing the rules alongside the measurements shows how the rules were applied.

We tested this approach with lymphocyte activation gene 3 (LAG-3), an immune checkpoint receptor whose ligands include major histocompatibility complex class II (19, 20) and fibrinogen-like protein 1 (21). LAG-3 was initially implicated in the transmission of α-synuclein preformed fibrils (22), but subsequent work challenged this role (23). A later study implicated LAG-3 in the transmission of pathological tau (24). In this campaign, we sought compounds that disrupted LAG-3 interactions with tau and amyloid-β while preserving binding to major histocompatibility complex class II and fibrinogen-like protein 1. Requiring both disruption and preservation provided a way to distinguish compounds with the desired profile from those that broadly interfered with LAG-3 interactions. Tau and amyloid-β are major therapeutic targets in Alzheimer’s disease (25, 26), and the PS19 (27) and 5xFAD (28) mouse models provide complementary settings for evaluating tau and amyloid pathology. To test whether the screening rules could transfer beyond LAG-3, we selected two cell-surface proteins linked to distinct aspects of neurodegeneration. Cellular prion protein (PrPᶜ) has been reported to mediate effects of amyloid-β oligomers on synaptic plasticity (29), and low-density lipoprotein receptor-related protein 1 (LRP1) regulates tau uptake and spread (30).

In our previous work, HTS-Oracle demonstrated that machine learning could prioritize compounds for prospective experimental testing across multiple targets. Subsequent studies applied this approach to immune checkpoint binders and extended validation to patient-derived cells (31–33). We asked whether an AI agent could use a specified pattern of interactions to guide an entire discovery campaign. In this setting, evidence that a compound disrupts one interaction is only part of the decision. Advancement also depends on showing that the interactions designated for preservation remain intact. Here we present Bivexia, an AI agent that guides a multistage pipeline from hit discovery and de novo design through preclinical evaluation. Its prespecified decision rules govern which compounds are selected and advanced against a prespecified interaction profile. We demonstrated the full pipeline on LAG-3 and tested the same screening rules prospectively on PrP^c^ and LRP1. We also report an exploratory comparison with medicinal chemist selections and examine where prospective screening outcomes diverged from retrospective predictions.

## RESULTS

### Bivexia guided selection of the target and interaction profile

Bivexia began with a planning stage to identify a tractable discovery problem involving extracellular protein interactions relevant to neurodegeneration. Given this domain, it proposed a target and specified which interactions to modulate and which to preserve. The planning prompt left the number of interactions in each group open and required comparison of candidate targets, explicit rejection criteria, and a distinction between experimental evidence and assumptions.

Bivexia compared five receptors and proposed LAG-3 as the initial target for testing whether small molecules could disrupt selected interactions while preserving others. PrPᶜ and LRP1 were reserved for subsequent tests of transfer across targets. For PrPᶜ, the aggregation state of its ligands required careful control, whereas domain-specific binding and trafficking could complicate the interpretation of initial LRP1 results (29, 30). In its planning record, Bivexia deprioritized TREM2 on the grounds that a purified interaction panel would capture only part of its context-dependent biology. It also excluded CD33 because its principal ligand recognition is glycan-based.

For LAG-3, Bivexia proposed disrupting interactions with tau and amyloid-β while preserving binding to MHC class II and FGL1. The preservation assays made selectivity an explicit requirement by testing whether compounds retained the other receptor interactions. This requirement shaped the objective function used throughout the campaign. Its preservation term used the lower of the two counterscreen scores, ensuring that a favorable prediction for one partner could not compensate for poor preservation of the other.

### A four-assay model selected compounds for a prespecified LAG-3 cascade

We first applied Bivexia to LAG-3, using a biochemical screening matrix of 8,960 compounds measured against four interactions: LAG-3 with tau, amyloid-β, MHC class II, and FGL1. In this matrix, 263 compounds (2.94%) met the criteria for disrupting the tau and amyloid-β interactions while preserving the MHC class II and FGL1 interactions. Bivexia used molecular fingerprints and physicochemical descriptors to predict these four outcomes for untested compounds. Its ranking function rewarded predicted disruption of tau and amyloid-β binding while limiting the score by the lower of the two predicted preservation probabilities.

Bivexia selected compounds for testing in successive batches and retrained its model on returned results. Thus, the model selected what to test; the policy determined what passed each experimental gate. Before the first assay, we committed a seven-stage advancement policy specifying the measurements, criteria for advancement, and conditions for retesting or stopping a compound (Fig. 1b,c and Table 1). The cascade progressed from primary biochemical screening through orthogonal binding and interference assays, cellular and Alzheimer’s disease donor-derived assays, receptor-dependence testing, pharmacokinetic and brain-exposure assessment, and efficacy studies in PS19 and 5xFAD mice. Table 1 summarizes the criteria and compound progression.

**Fig. 1.**
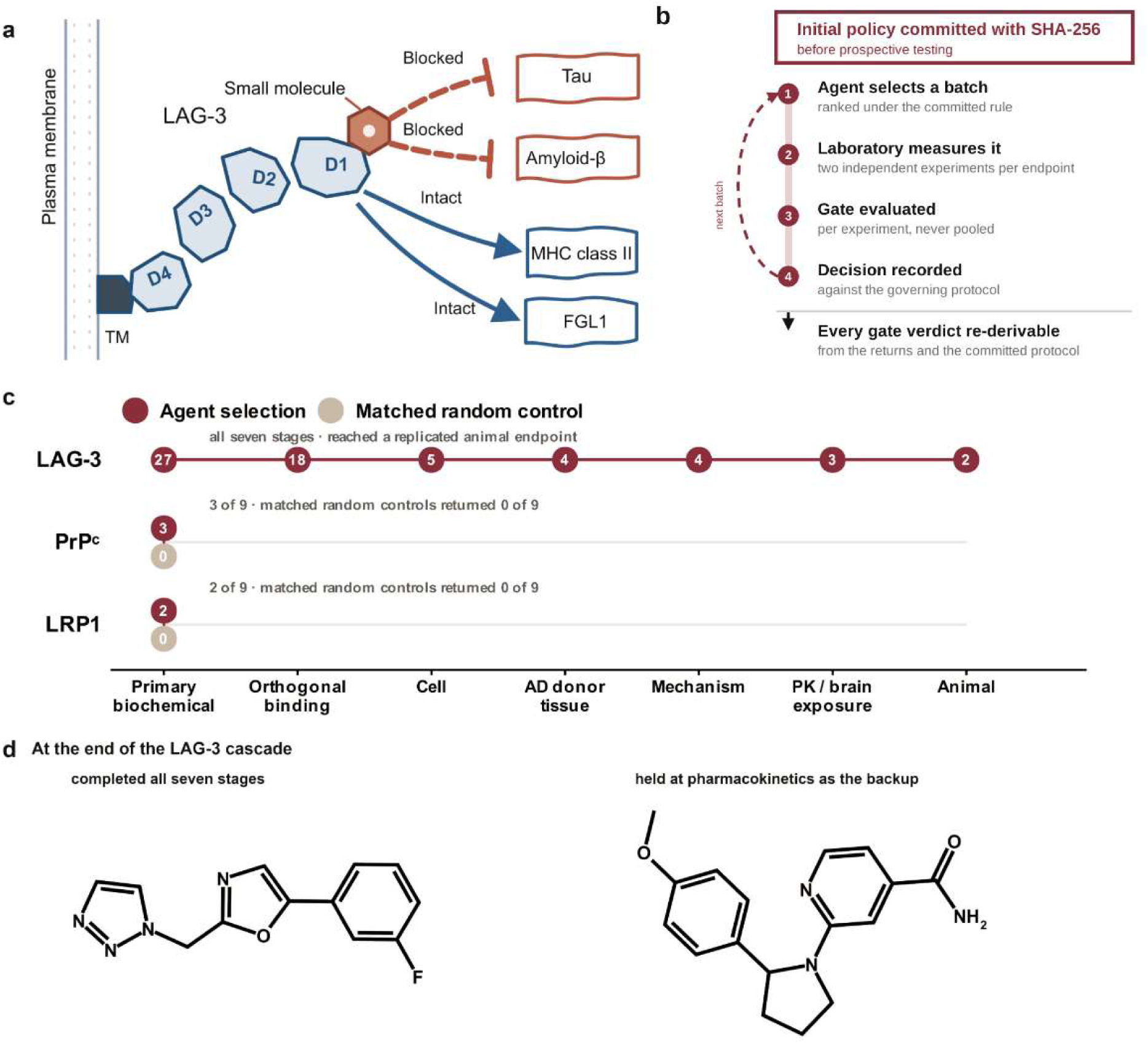
Bivexia and the prespecified decision policy. **a**, Discovery objective. A compound must block the LAG-3 interactions with tau and amyloid-β while preserving those with MHC class II and FGL1. **b**, Experimental workflow. The initial policy was committed before prospective testing. Gate verdicts are evaluated per independent experiment and are re-derivable from the returns and protocol. **c**, Stage counts for LAG-3 and primary-screen outcomes for PrPᶜ and LRP1. The LAG-3 animal count records two logged entries; one compound was dosed, and the backup promotion was reverted before any animals were committed. **d**, Structures of the completed lead (BVX-01) and the backup (BVX-02), which remains held at the pharmacokinetic stage.

**Table 1.**
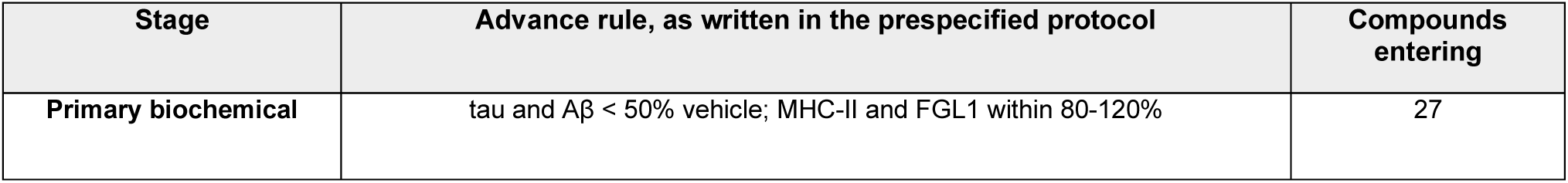
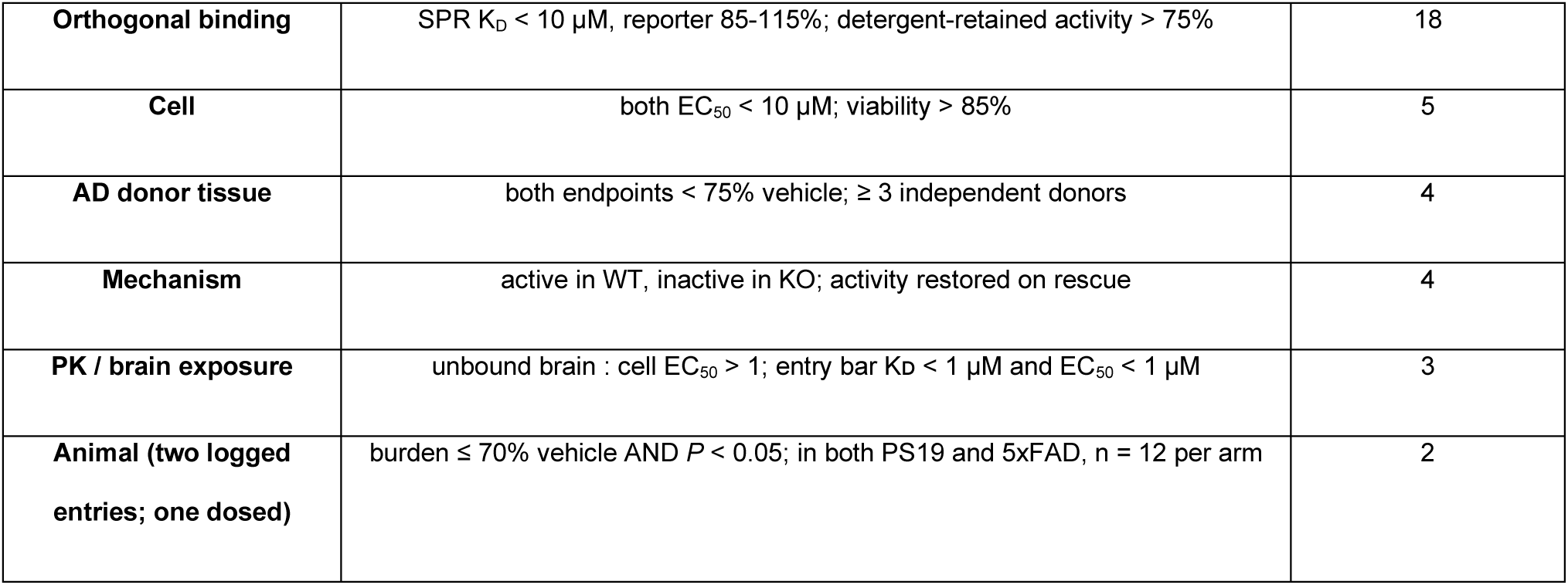
Prespecified advancement criteria and compound progression. Criteria are summarized from the initially committed protocol.

| Stage | Advance rule, as written in the prespecified protocol | Compounds entering |
|---|---|---|
| **Primary biochemical** | tau and Aβ < 50% vehicle; MHC-II and FGL1 within 80-120% | 27 |
| **Orthogonal binding** | SPR K<sub>D</sub> < 10 µM, reporter 85-115%; detergent-retained activity > 75% | 18 |
| **Cell** | both EC<sub>50</sub> < 10 µM; viability > 85% | 5 |
| **AD donor tissue** | both endpoints < 75% vehicle; ≥ 3 independent donors | 4 |
| **Mechanism** | active in WT, inactive in KO; activity restored on rescue | 4 |
| **PK / brain exposure** | unbound brain : cell EC<sub>50</sub> > 1; entry bar Kᴅ < 1 µM and EC<sub>50</sub> < 1 µM | 3 |
| **Animal (two logged entries; one dosed)** | burden ≤ 70% vehicle AND *P* < 0.05; in both PS19 and 5xFAD, n = 12 per arm | 2 |

Across the campaign, 27 compounds entered the cascade. Eighteen passed the primary screen (fig. S1), five passed orthogonal binding and interference counterscreens, and four advanced through the Alzheimer’s disease donor-derived assays. BVX-01 completed the cascade; BVX-02 met the pharmacokinetic criteria and was retained as a backup (Fig. 1c,d and Table 1).

### BVX-01 completed the discovery cascade and reduced pathology in two mouse models

BVX-01 bound human LAG-3 with a dissociation constant of 42 ± 4.0 nM and mouse LAG-3 with a dissociation constant of 135 ± 9.2 nM. We then tested its functional activity in two cell assays: tau seeding in Tau RD P301S CFP/YFP biosensor cells, and Aβ42 uptake in primary mouse cortical neurons. Across two independent experiments, the fitted EC₅₀ values were 0.22 and 0.25 μM for tau seeding and 0.28 and 0.32 μM for Aβ42 uptake (fig. S2). At 10 μM, cell viability remained 92.8% of vehicle.

We next tested BVX-01 in induced cortical neurons from three independent donors with Alzheimer’s disease. At 10 μM, tau-seeding signals were 30.1%, 37.5%, and 39.6% of vehicle across the three donors; Aβ42-association signals were 49.0%, 55.7%, and 59.9%. Each donor met the prespecified criterion of less than 75% of vehicle. In matched LAG-3-knockout cells, the tau-seeding and Aβ42-uptake responses returned to 93.5% and 94.8% of vehicle, consistent with the compound’s effects requiring LAG-3.

Following oral administration, peak unbound brain concentrations exceeded the cellular tau-seeding EC_50_ by factors of 2.41 and 2.82, meeting the exposure criterion for advancement. We then tested whether BVX-01 reduced pathology-associated brain readouts in two complementary mouse models. Daily oral treatment reduced hippocampal tau-aggregation signal to 60.0% of vehicle in PS19 mice and brain Aβ42 levels to 63.6% in 5xFAD mice. Independent confirmatory cohorts reproduced the reductions, yielding corresponding values of 52.9% and 59.2%. All four comparisons met the prespecified efficacy criterion of no more than 70% of vehicle with *P* < 0.05 (Fig. 2a,b). A brain thermal-shift assay provided additional evidence consistent with LAG-3 engagement (Fig. 2c). Body weight remained at or above 90.3% of baseline, meeting the prespecified weight-retention criterion (Fig. 2d).

**Fig. 2.**
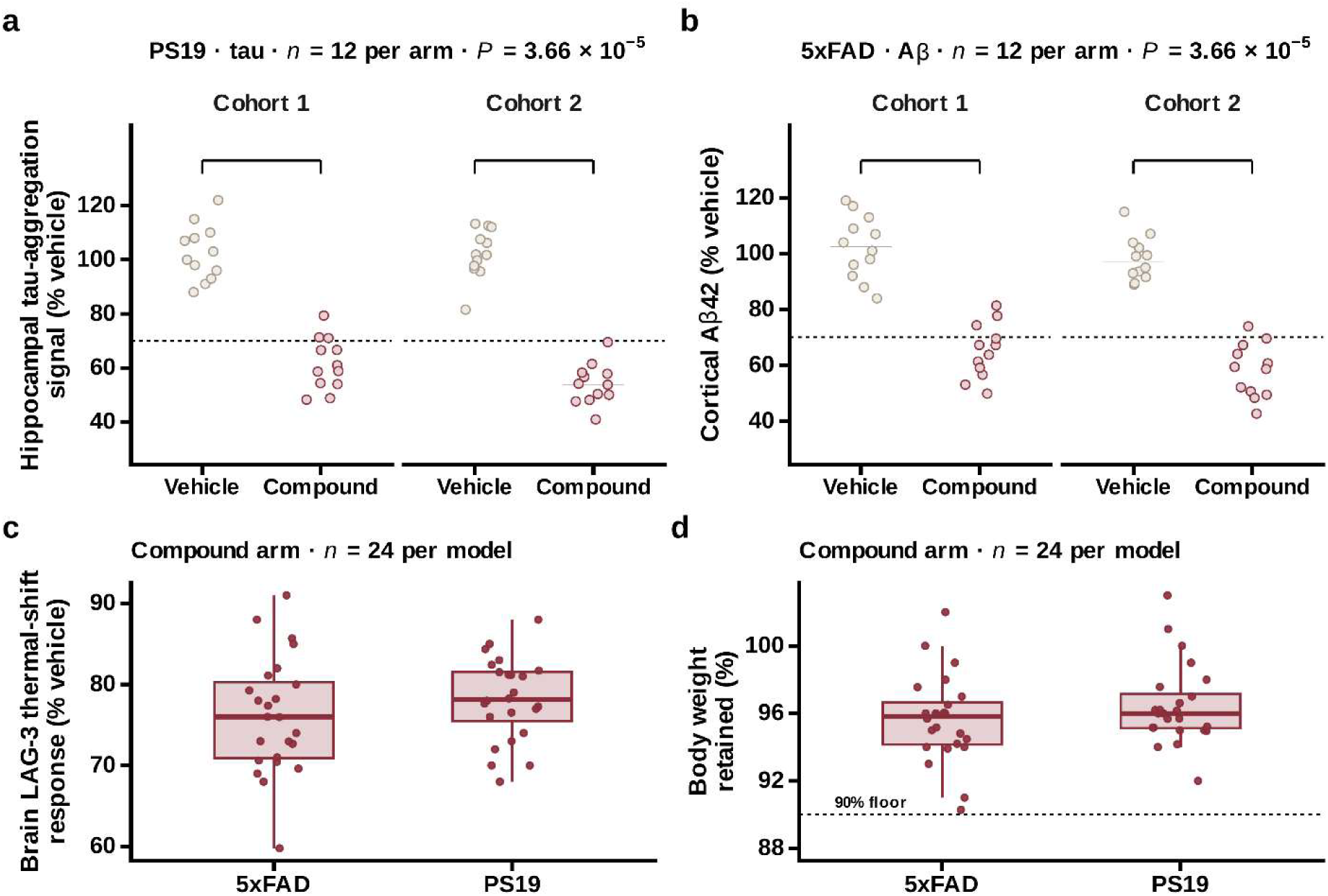
In vivo efficacy and tolerability of BVX-01. a, Tau burden in PS19 mice represented by hippocampal tau aggregation and **b,** Aβ burden in 5xFAD mice represented by cortical Aβ, shown for two independent cohorts. Every animal is plotted (n = 12 per arm); the horizontal rule is the group median and the dashed line marks the prespecified 70% advance threshold. Brackets denote vehicle versus compound. Vehicle and compound distributions do not overlap in any of the four comparisons. The runner used a two-sided Mann-Whitney *U* test with the asymptotic approximation. **c,** Brain LAG-3 thermal shift response (%vehicle) and **d,** body weight retained, in treated arms of both models; the dashed line in d marks the 90% tolerability bound. Thresholds and sample sizes were set in the initially committed protocol.

BVX-01 retained the interaction selectivity that motivated its advancement. In separate biochemical assays, residual LAG-3 binding to tau and amyloid-β was approximately 35-45% of vehicle, whereas binding to MHC class II and FGL1 remained at 90-105%. Thus, inhibition of the designated interactions occurred without substantial loss of binding to the preserved partners under the tested conditions. This functional separation prompted us to investigate which receptor regions supported compound binding and inhibition while retaining the preserved interactions.

### De novo design yielded selective inhibitors and tested a pharmacophore hypothesis

We next asked whether Bivexia could extend beyond catalogue selection to design new compounds with the desired interaction profile. We enumerated products accessible through a single amide coupling of commercially available building blocks, linking molecular design to a practical synthesis route. Nine compounds (fig. S3) were selected, and their structures were cryptographically committed before synthesis. The commitment specified a success criterion of at least two primary hits among nine compounds, together with a one-sided exact binomial test, a library base rate of 0.029, and the corresponding probability table.

Five of the nine compounds met the primary screening criteria (Fig. 3a). Under the committed base rate, the expected count was 0.26, and the probability of observing five or more hits was 2.3 × 10^-6^ (Fig. 3b). Using the more precise measured library rate of 2.94% yielded a similar probability of 2.5 × 10^-6^. All five primary hits were carried to orthogonal confirmation. DN-05, DN-01, and DN-02 confirmed, with SPR dissociation constants for LAG-3 binding of 78, 155 and 210 nM, respectively (Fig. 3c). Of the other two, one bound at 300 nM but fell outside the committed 85-115% reporter window at 122%, indicating assay interference, and the other bound above the 10 µM threshold. The remaining four compounds were also profiled, to characterize the designed series as a whole: the two triazole-free controls described below, and two further compounds, one of which was stopped for colloidal aggregation at 72% detergent-retained activity against a 75% requirement.

**Fig. 3.**
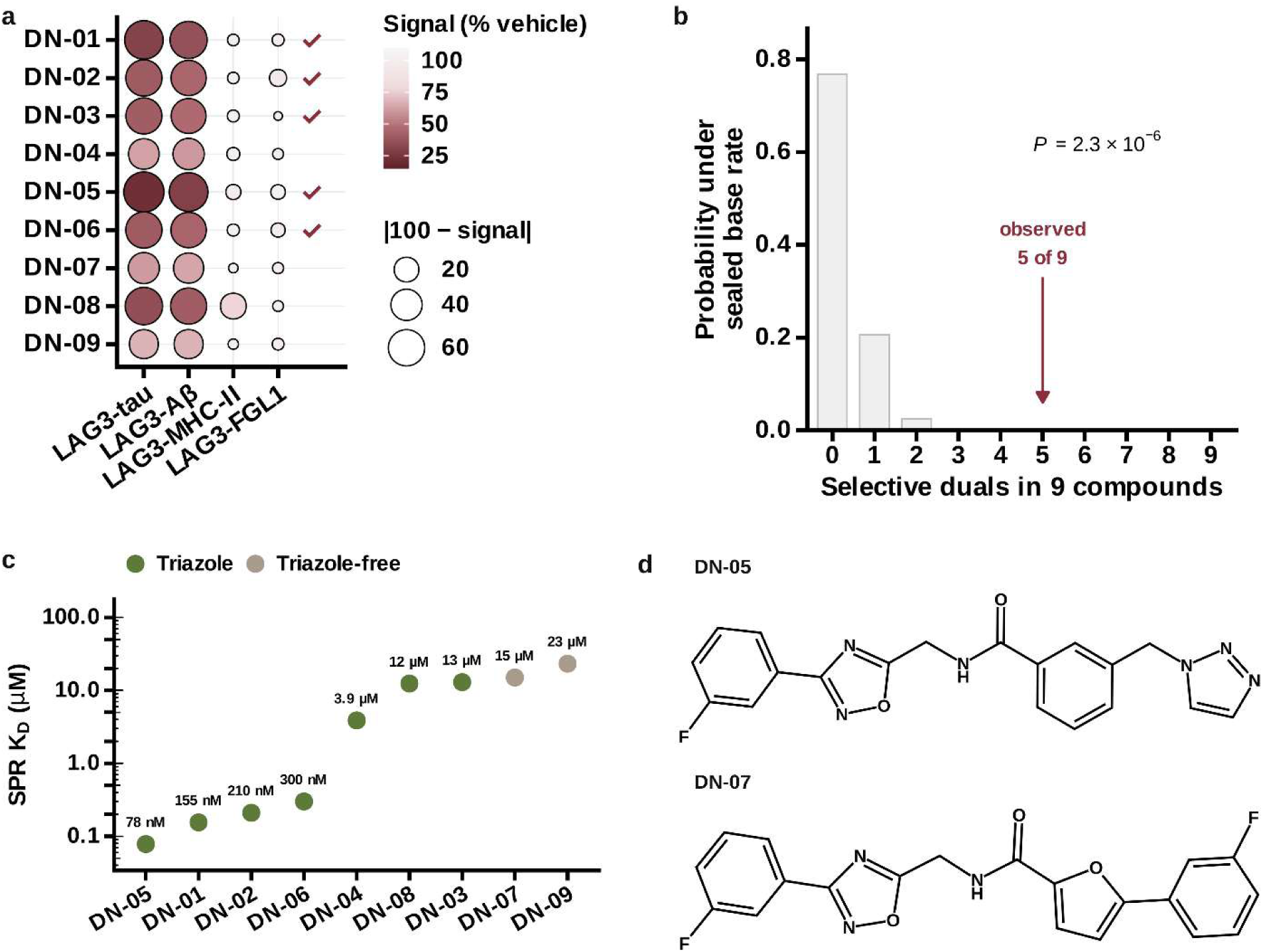
De novo design using Bivexia. **a**, Four-assay profile of all nine compounds. Point position is the mean of independent experiments, fill is signal as a percentage of vehicle and size is deviation from vehicle; a filled marker at the right indicates a compound meeting all four criteria in every independent experiment. **b**, The observed count against the null distribution implied by the base rate of 0.029; the expected count is 0.26 and *P* = 2.3 × 10^-6^ by the one-sided exact binomial test committed before synthesis. The measured library rate of 2.94% gives 2.5 × 10^-6^. **c**, Orthogonal SPR affinities for all nine compounds, ordered by affinity and colored by chemotype. **d**, The tightest binder (DN-05) and a triazole-free member of the same series (DN-07), which together test the pharmacophore hypothesis; their affinities and chemotypes are given in **c**. Structures of all nine compounds are given in fig. S3 and building-block identifiers in table S2.

The series also incorporated a test of the proposed pharmacophore. Two compounds lacking the triazole element (DN-07 and DN-09) showed substantially weaker binding, with dissociation constants of 15 and 23 µM, compared with 78 nM for the strongest binder DN-05 (Fig. 3c,d and fig. S4). Their inclusion in the commitment before synthesis provided prospective controls for the contribution of this element. These results support a role for the triazole-containing motif in binding affinity. We subsequently examined whether the weaker binding of a triazole-free compound was accompanied by altered dependence on the receptor region implicated by deletion mapping.

### Two chemotypes share a regional dependence within LAG-3 domain 1

We next asked whether the shared selectivity profile of the catalogue-selected lead (BVX-01) and the strongest de novo binder (DN-05) reflected a common dependence on a region of LAG-3. We compared their binding and activity across three deletion constructs spanning domain 1 and seven point mutants at residues implicated in interactions with MHC class II or FGL1 (19–21). The panel was tested against the campaign lead (BVX-01), DN-05, and the weaker triazole-free compound DN-07 (Fig. 4a).

**Fig. 4.**
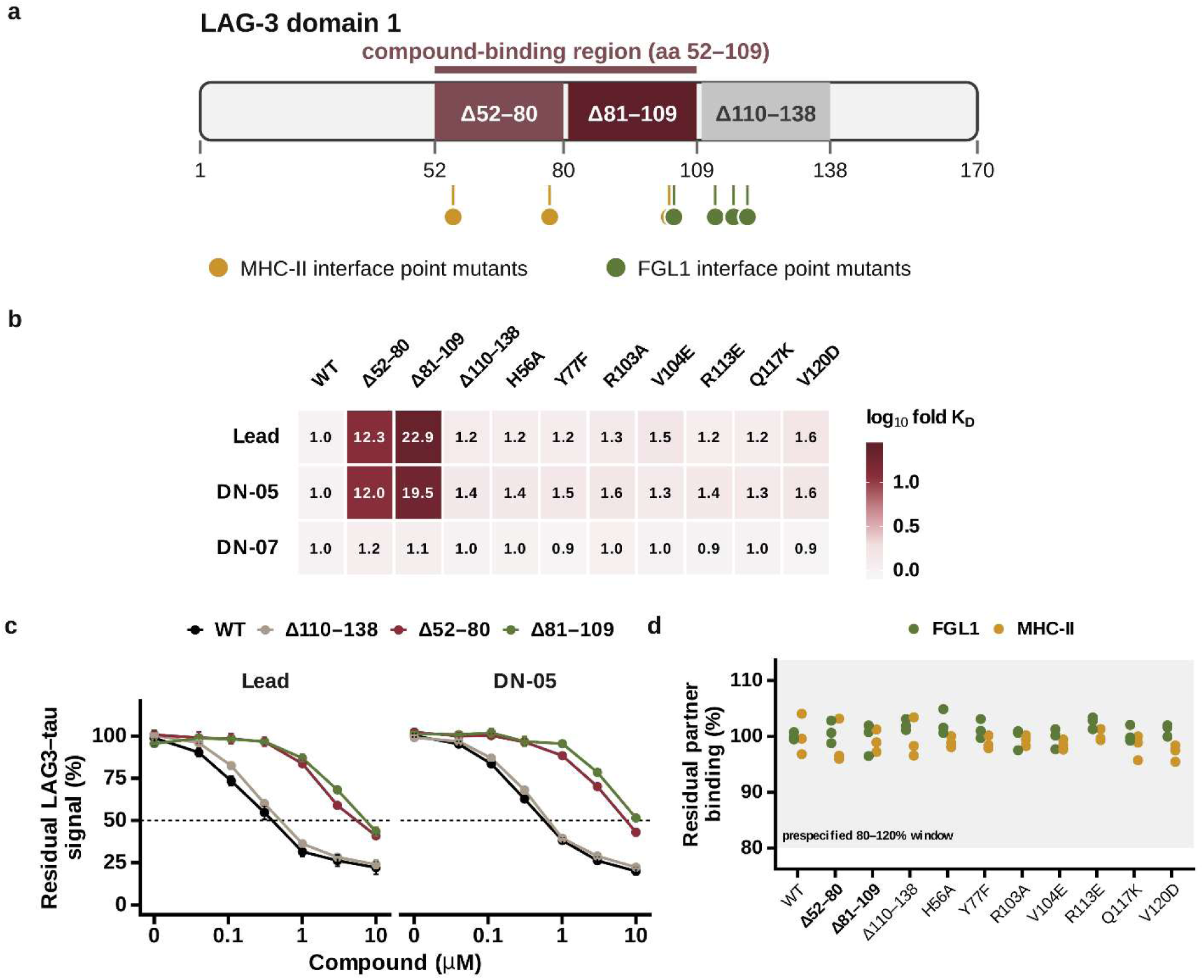
Deletion mapping places both chemotypes on LAG-3 D1 residues 52-109. **a**, Constructs tested. Three deletions span domain 1; seven point mutants mark residues at the MHC class II and FGL1 interfaces. **b**, Fold change in dissociation constant relative to wild type for each compound and construct; numbers are fold change and color is its base-10 logarithm. The underlying per-preparation dissociation constants are given in fig. S5. **c**, Dose-response for residual LAG3-tau signal; points are means of three independent preparations and error bars are s.e.m. **d**, Preserved-partner binding in the presence of compound across all constructs. Every protein preparation is plotted and the shaded band is the prespecified 80-120% acceptance window; the two deletions spanning 52-109, which lose compound binding, are set in bold on the axis. Basal partner binding without compound is shown in fig. S6a, and the corresponding amyloid-β concentration-response curves are shown in fig. S6b.

The two active compounds (BVX-01 and DN-05) showed closely matched responses to the deletion panel. For BVX-01, dissociation constants increased 12.3-fold on Δ52-80 and 22.9-fold on Δ81-109. The adjacent deletion, Δ110-138, produced a 1.2-fold change, while the seven point mutants produced changes of 1.2-1.6-fold (Fig. 4b,c). DN-05 showed corresponding increases of 12.0-fold and 19.5-fold on the two deletions spanning residues 52-109, compared with 1.3-1.6-fold elsewhere (Fig. 4b and fig. S5). DN-07 showed little sensitivity to any construct, with changes ranging from 0.9-fold to 1.2-fold across the full panel (Fig. 4c and fig. S5). The functional measurements followed the same pattern. At 10 µM, both active compounds reduced the residual LAG-3-tau interaction signal to 22-24% of vehicle on wild-type LAG-3 and the point mutants. Residual signals increased to 41-52% on Δ52-80 and Δ81-109, accompanied by shifts toward higher concentrations in the dose-response curves (Fig. 4c). The corresponding amyloid-β results are shown in fig. S6b. DN-07 produced weaker effects throughout the panel, with residual signals of 70-77%.

We examined protein integrity and basal partner binding to assess whether deletion effects could reflect broader changes in receptor function. The two deletions spanning residues 52-109 reduced melting temperatures by approximately 3 °C and retained 86-89% monomer by size-exclusion chromatography, compared with 96% for wild-type LAG-3. Both constructs retained approximately 80% of basal tau and amyloid-β binding (fig. S6a). The point mutants showed integrity measurements comparable to wild type, and each reduced binding of its intended partner, demonstrating sensitivity to perturbations at the preserved-partner interfaces. In the presence of compound, MHC class II and FGL1 binding remained at 97-105% of the corresponding control across the construct panel (Fig. 4d).

Our results implicate residues 52-109 in the binding and activity of both chemotypes. The retained monomer content and basal partner binding argue against gross loss of receptor integrity, although the modest destabilization leaves open a contribution from local conformational changes. Resolving direct contacts within this 58-residue region will require finer mapping. Bivexia selected the catalogue compound BVX-01 and designed DN-05 using an objective based on four assay outcomes, without an explicit binding-site constraint. Their shared deletion sensitivity therefore connects the prospectively selected interaction profile to a common regional dependence within LAG-3. The weaker triazole-free compound lacked this pattern, further distinguishing its binding behavior from that of the two active chemotypes.

### The screening policy yielded hits on two additional receptors

We next tested whether the decision rules established for LAG-3 could support prospective discovery on PrPᶜ and LRP1, two receptors implicated in the handling of pathological aggregates (29, 30). Both had been nominated as cross-target tests during the initial planning stage. We generated their protocols programmatically from the LAG-3 protocol. After endpoint renaming, all gates, thresholds, boundary bands, sample sizes, retest limits, and stopping rules were identical. The only recalculated parameter was the batch success criterion, which depends on the measured hit rate for each target. This design allowed us to test transfer without retuning the advancement rules.

The screening matrices contained substantially fewer selective inhibitors for the additional receptors. Before prospective selection, 25 of 8,960 compounds met the criteria for PrPᶜ (0.279%) and 12 of 8,960 for LRP1 (0.134%), compared with 263 of 8,960 for LAG-3 (2.94%). At these lower base rates, one hit among nine compounds would meet the probability threshold used to derive the LAG-3 criterion. We nevertheless retained the stricter requirement of at least two hits among nine compounds for both targets.

Retrospective evaluation provided little evidence that the agent would outperform random selection. Under scaffold-grouped five-fold cross-validation using the labels for each target, top-k enrichment was approximately onefold for PrPᶜ and LRP1, compared with 17.7-fold for LAG-3 under the same procedure (Fig. 5a). Retrospective sequential campaigns with retraining across 990 compounds likewise showed no separation from random selection on either target (fig. S7). We recorded this limited support in each prospective batch commitment and specified that a null outcome would also be reported.

**Fig. 5.**
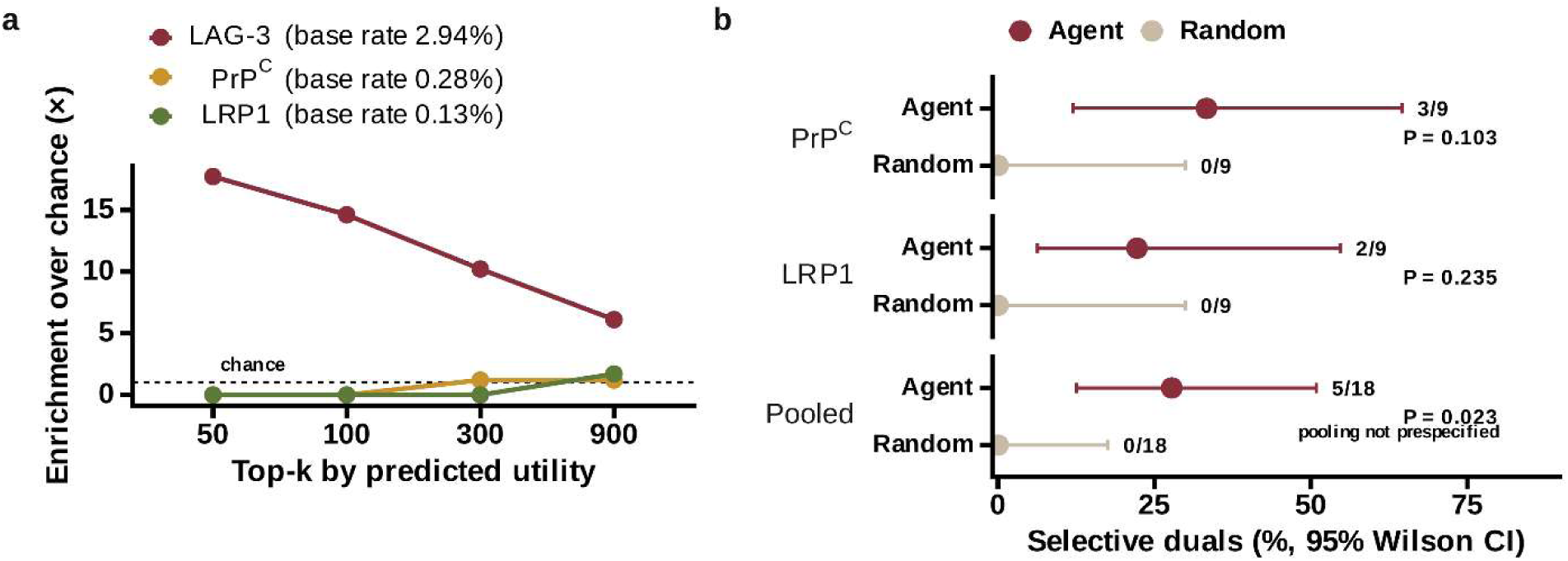
Transfer of the prespecified policy to PrPᶜ and LRP1. **a**, Top-k enrichment over chance under scaffold-grouped five-fold cross-validation on each target’s own labels, with each target’s measured library base rate given in the legend; the ported targets show essentially no enrichment where LAG-3 shows 17.7-fold at top-50. **b**, Hit rates with 95% Wilson confidence intervals, agent against the matched random control drawn from the same pool, for each target and pooled; counts are given beside each interval and *P* values are one-sided Fisher’s exact tests. Neither target reaches significance alone at n = 9 per arm. The pooled result is a post hoc, unadjusted summary and is not confirmatory. Retrospective enrichment did not predict prospective performance: both ported targets returned selective inhibitors well above their library base rates despite showing no cross-validation enrichment. Sequential campaigns underlying **a** are shown in fig. S7, and the near-neighbor control excluding retrieval of known actives in fig. S8.

For each target, the agent ranked 1,675 candidate structures without recorded measurements in any assay matrix, applied the same applicability-domain filter, and selected nine compounds. A matched random batch of nine compounds was drawn from the same eligible pool. All four selection lists were cryptographically committed before screening. The random-control commitments also specified how the observed counts would be interpreted, although the small batches limited how precisely they could characterize the underlying pool hit rates.

Prospective screening yielded three hits among nine agent selections for PrPᶜ and two among nine for LRP1. Neither random batch yielded a hit (Fig. 5b). Both agent batches therefore met the prespecified criterion of at least two hits, although the planned comparisons against their respective random controls did not reach conventional significance (one-sided Fisher exact *P* = 0.103 and *P* = 0.235). In an exploratory, post hoc summary across the two targets, 5 of 18 Bivexia-selected compounds and 0 of 18 matched random selections met the hit criteria (nominal, unadjusted one-sided Fisher’s exact *P* = 0.023). Because this pooled analysis was not prespecified, we treat it as descriptive, not as confirmatory evidence that Bivexia outperforms random selection. The individual assay profiles showed why the preservation criteria mattered. One random-control compound met both disruption criteria on each target but failed the preservation criteria, demonstrating that disruption alone was insufficient to produce a selective hit. A compound identified as a PrPᶜ hit also appeared by chance in the LRP1 random batch, where it failed the disruption criteria. This difference indicates variation in activity between the two assay panels, although broader cross-panel profiling was not prespecified.

The prospective hit rates corresponded to 119-fold and 166-fold enrichment over the measured library base rates for PrPᶜ and LRP1, respectively. These values provide historical context, because the prospective candidates came from a separate pool whose hit rate was estimated only through the small random batches. Similarity analysis also offered limited support for retrieval of close analogs of known actives. Maximum Tanimoto similarities between prospective hits and training-set selective inhibitors ranged from 0.20 to 0.53. Hits were no closer to training actives than misses (fig. S8), and none shared a Bemis-Murcko scaffold (34) with a training active. The difference between retrospective enrichment and prospective outcomes therefore remains unresolved. The results establish that the unchanged screening rules yielded compounds meeting the interaction criteria on two additional receptors, while larger prospective comparisons are needed to assess the selection advantage on each target.

### An exploratory comparison with outcome-blinded medicinal chemists

We next examined how compound selection by Bivexia compared with selections made independently by two medicinal chemists who were blinded to assay outcomes. Four arms each selected 30 compounds. The agent and seeded random selections were cryptographically committed before screening, and both lists verify against their recorded digests. Selections were allocated sequentially. Removing compounds with an existing campaign status left 805 candidates without available assay outcomes. Bivexia selected 30 compounds deterministically, and a seeded random draw selected 30 from the remaining 775. Both chemists then independently selected from the 745 compounds left after these allocations. All four selection sets were non-overlapping, and all 120 compounds were screened together using the same protocol. This sequence gave Bivexia first access to the candidate pool and prevented either chemist from selecting compounds already assigned to the agent or random arm.

We examined the utility distribution to characterize the candidates that remained available to the chemists (Fig. 6a). Forty-four of the 745 compounds scored at or above the lowest utility among the agent selections. The highest-scoring compound in the original pool also remained available, with a utility of 0.62 compared with 0.48 for the highest-scoring agent selection. Median utilities for the two chemist selections were 0.212 and 0.204, close to the pool median of 0.201. The agent selections averaged approximately the 95th percentile relative to that same 745-compound pool. These distributions show that high-scoring candidates remained available after the initial allocations, while leaving the effect of selection order unresolved.

**Fig. 6.**
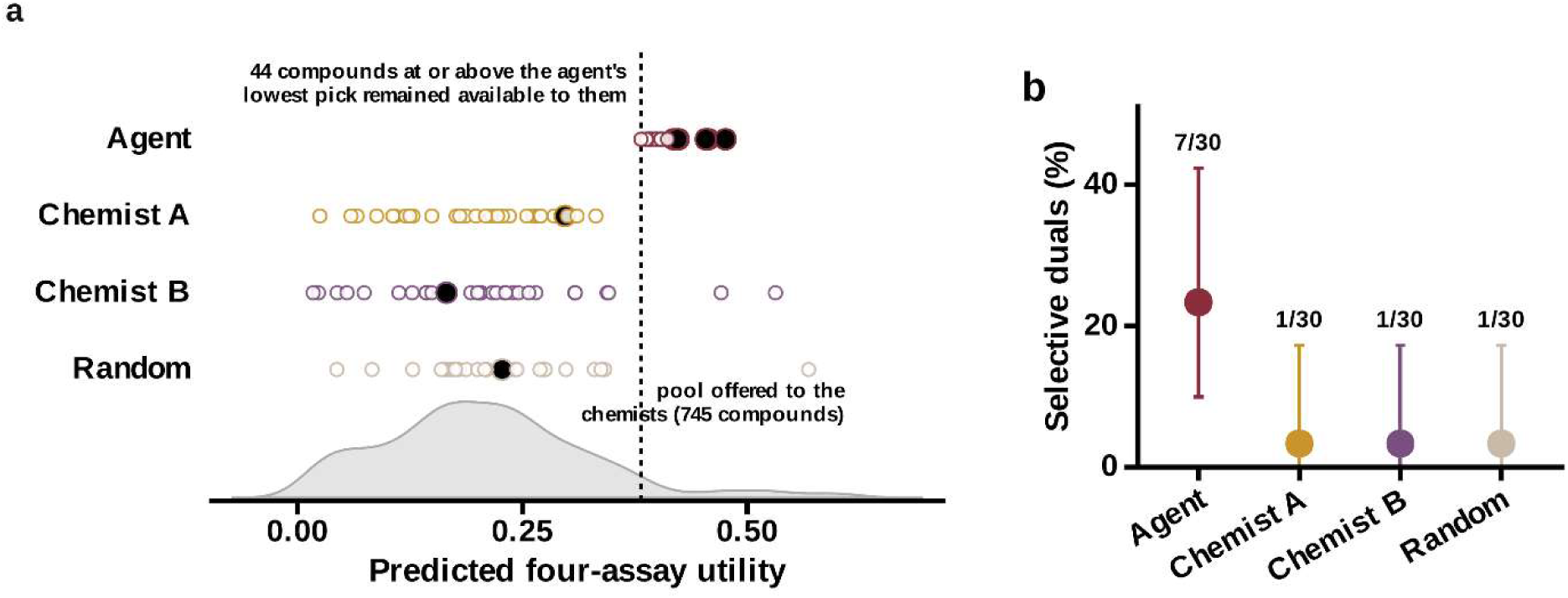
Supporting comparison against medicinal chemists. a, Every committed compound on the model’s own utility scale. Each circle is one of 30 compounds committed by hash per arm; filled circles are selective inhibitors (disruption of both aggregate interactions below 50% of vehicle with both preserved interactions held within 80-120%, evaluated in every independent experiment). The shaded curve is the distribution of predicted four-assay utility across the 745 compounds offered to the chemists. Selection was sequential: the agent drew 30 from 805, the seeded random arm drew 30 from the remaining 775, and the chemists then selected from the 745 that remained, so no arm could take a compound another had taken. The dashed line marks the agent’s lowest-ranked selection: 44 compounds scoring at or above it, including the highest-scoring compound in the set, remained available to the chemists, whose selections sat near the pool median. Utility values are from the model as fitted during the campaign. **b,** Measured hit rate for all four arms with 95% Clopper-Pearson intervals and counts. Both chemists recovered selective inhibitors at the same rate as random selection.

Bivexia identified seven selective inhibitors among 30 selections, compared with one among 30 for each chemist and one among 30 for random selection (Fig. 6a,b). The primary hits from all four arms were carried forward to orthogonal confirmation under the same gates used in the main campaign. Six of the seven Bivexia hits confirmed, and none of the three comparator-arm hits did: six confirmed binders per thirty committed compounds for Bivexia (95% CI 7.7-38.6%), against none per thirty in each comparator arm (95% CI 0-11.6%). The higher observed yield warrants further evaluation under a design that gives all selectors access to the same candidate pool. Such a comparison would require outcome-blinded selections with the selection lists, primary comparison, and analysis plan committed before screening.

## DISCUSSION

Bivexia demonstrates how an AI agent can guide a discovery campaign toward a specified pattern of protein interactions. Compound selection, de novo design, and advancement were organized around a common objective of disrupting selected interactions while preserving others. Bivexia links an explicit interaction-selectivity objective to compound selection, de novo design, and experimental advancement. In the LAG-3 campaign, BVX-01 completed a seven-stage cascade and reduced tau and amyloid-β pathology in two mouse models, with effects reproduced in independent cohorts. The de novo series produced five primary hits among nine compounds, three of which bound LAG-3 with nanomolar affinity. Applying the screening rules to PrPᶜ and LRP1 yielded three and two selective hits, respectively, among nine agent-selected compounds per target; neither matched random batch produced a hit. Together, these results extend prospective interaction-selective discovery beyond a single receptor while demonstrating the full preclinical cascade for LAG-3.

The LAG-3 campaign shows how preservation requirements can shape discovery. BVX-01 disrupted the tau and amyloid-β interactions while retaining binding to MHC class II and FGL1. These preservation outcomes directly constrained the compound-ranking objective, so compounds could not rank highly by disrupting the designated interactions while broadly impairing LAG-3 binding. The objective therefore guided selection toward a defined interaction profile rather than general inhibition of the receptor.

Binding-site mapping connected this functional profile to a shared regional dependence. BVX-01 and the de novo compound DN-05 lost affinity on deletions spanning LAG-3 residues 52-109, whereas the weaker triazole-free compound DN-07 showed little sensitivity to the same deletions. The selection objective contained no binding-site constraint, so the shared pattern emerged from selection for the four assay outcomes. These experiments identify a region associated with binding and activity, not direct molecular contacts; local structural effects of the deletions remain possible.

Retrospective analyses also shaped Bivexia’s selection strategy. We used exploit-only batches with retraining in the prospective campaign. The utility score should be interpreted within its intended task: it prioritizes compounds for the specified interaction profile, and its rank relationship to measured affinity is not stable across chemical sets (35). Direct assays remain necessary to establish affinity and potency.

On PrPᶜ and LRP1, the prospective batches met their prespecified hit criteria despite showing little enrichment in scaffold-grouped cross-validation. These outcomes demonstrate that Bivexia identified compounds meeting the intended interaction profiles on two additional receptors. The small target-specific comparisons and exploratory pooled analysis limit how precisely this study can estimate performance relative to random selection; the pooled analysis and individual comparisons are reported in the Results. The exploratory medicinal chemist comparison yielded seven selective inhibitors among 30 Bivexia selections, compared with one among 30 for each chemist and one among 30 for random selection. Carrying those primary hits through the campaign’s orthogonal gates separated the arms further: six of the seven Bivexia hits confirmed as direct LAG-3 binders, whereas none of the three comparator-arm hits did. Measured as confirmed binders recovered per thirty committed compounds, the endpoint a discovery campaign ultimately depends on, the comparison is six against none. Each comparator arm contributed only a single primary hit, so this separation follows mainly from the difference at the primary screen rather than from a measured difference in hit quality. The result motivates a more controlled benchmark, but the arms did not have equivalent access to candidates: Bivexia selected first, and the chemist lists were hashed after screening.

Several limitations define the next steps. The full cascade rests on one compound, and the PrPᶜ and LRP1 campaigns stopped at primary screening. The biological role of the LAG-3-amyloid-β interaction is less established than that of LAG-3-tau, and the mouse results do not establish how much each disrupted interaction contributed to the reductions in pathology. Further studies across compounds, targets, and partner panels will test the breadth of the approach. Bivexia provides a framework for carrying a defined interaction objective through prospective decisions, with protocols, code, and measurements available for evaluation and extension.

## MATERIALS AND METHODS

### Decision policy and advancement rules

The initial decision policy was a JSON document specifying the seven stages, endpoints, thresholds, retest behavior, sample sizes, campaign-level halt rules, proposal throttle, and animal-use ceiling. Every endpoint is evaluated per independent experiment, and a compound advances only if the criterion is met in all of them. Six amendments were subsequently recorded, each superseding a named protocol digest. The six amendments added decision machinery absent from the initial document; none altered a gate, threshold, sample size, or statistical test. Technical replicates are averaged within an experiment. Applying the per-experiment advancement rule did not change any primary-gate decisions, but it changed one orthogonal-stage decision. Detergent-retained activity was 71.9 and 72.2% in one experiment and 87.0 and 88.6% in the other. The pooled mean was 79.9%, above the >75% threshold, but the compound failed because it did not pass in each independent experiment. Primary advance requires both disrupt endpoints below 50% of vehicle and both preserved endpoints within 80 to 120%. Orthogonal advance requires a dissociation constant below 10 µM, reporter signal within 85 to 115%, and detergent-retained activity above 75%. In vivo statistics are computed by the runner from per-animal values using the two-sided Mann-Whitney U test with the asymptotic approximation at α = 0.05, with n = 12 per arm and dedicated vehicle arms in each model. Cellular advancement required tau-seeding and Aβ42-uptake EC_50_ values below 10 μM and viability above 85% in each independent experiment. Advancement from the Alzheimer’s disease donor-derived assays required both signals to be below 75% of matched vehicle in at least three independent donors. The receptor-dependence gate required activity in wild-type cells, loss of activity in LAG-3-knockout cells, and restoration of activity after LAG-3 rescue. Pharmacokinetic advancement required an unbound-brain-to-cellular-EC_50_ ratio greater than 1; entry into animal studies also required an SPR dissociation constant and cellular EC_50_ values below 1 μM. Animal advancement required the pathology endpoint to be no more than 70% of vehicle, with *P* < 0.05, separately in both models.

### Selection policy

A four-output model predicts the probability of a hit call on each endpoint. Each compound is represented by a 1024-bit Morgan count fingerprint of radius 2 (*36*) concatenated with ten physicochemical descriptors, giving 1,034 features; fingerprints and descriptors were computed with RDKit (*37*). One random forest classifier was fitted per endpoint with scikit-learn (*38*), using 400 trees, a minimum of four samples per leaf, the square root of the feature count considered at each split, balanced class weights, and a fixed random seed. The predicted probability is the mean vote across trees, and the standard deviation across trees is retained as a measure of model uncertainty. The exact feature specification is given in the Supplementary Methods and is recorded in the released code as the feature version string, which a saved model carries so that features and model cannot be mixed across versions. Compounds are ranked by the square root of the product of the two disrupt probabilities multiplied by the minimum of the two preserve probabilities. Similarity was computed separately from the model features, as the Tanimoto coefficient between 2048-bit binary Morgan fingerprints of radius 2. Candidates further than Tanimoto 0.40 from the nearest training compound were excluded as outside the applicability domain, as were candidates within Tanimoto 0.40 of any compound that failed a stage after the primary screen. The model is retrained on all returned results before each proposal. In the LAG-3 campaign, Bivexia selected compounds in three sequential batches of nine.

### Prospective commitments

The agent arm of the chemist comparison was committed more than once; each superseded commitment is retained in the repository with its digest and the reason it was replaced. The first was discarded because the candidate pool was found to contain compounds with existing assay outcomes, and was replaced after the pool was rebuilt from compounds carrying no campaign status; the second was replaced when the arm size was raised from nine to thirty, because at nine compounds against a 2.9% base rate no arm can distinguish selection skill from chance. The seeded random arm was not redrawn, since all of its members lay within the corrected pool and redrawing after the contamination had been observed would itself have been a selection informed by outcomes.

For every prospective arm the sorted compound identifiers were concatenated and hashed with SHA-256, and the digest written together with the selection rule, the pool, the criterion and the null. Table S1 lists each commitment with its payload definition, repository record, digest and timing. The de novo series, the ported-target batches and their random controls, and the agent and random arms of the chemist comparison were all hashed before any member was screened. The two chemist arms were hashed after screening; the investigators report that selection preceded screening, but no contemporaneous record substantiates that timing, so those digests fix which compounds were chosen and not when. Agent arms were generated deterministically by script from the pool file and the fitted model, and each commitment record stores the selection rule alongside the digest that fixes the compounds selected. Random arms are drawn from a declared seed with the recipe recorded. Chemist arms were committed independently by each chemist from a pool file containing compound identifier, SMILES and similarity to the training set, and no assay outcome.

### De novo enumeration

Products were enumerated by a single amide coupling between catalogue acid and amine building blocks, filtered for structural alerts and for novelty against the training set, and ranked by the same objective function. Candidate generation and scaffold analysis used RDKit (37); scaffolds are Bemis-Murcko frameworks (34).

### Cross-target porting

Ported protocols were generated by script from the LAG-3 protocol; the four endpoint names were substituted, the commitment criterion re-derived, and all other fields identical after renaming. Base rates were measured from each target’s assay matrix before any selection. The criterion was taken as the larger of the value derived at the false-positive probability accepted in the LAG-3 campaign and the source campaign’s value of 2 of 9. Both ported protocols were fixed before any compound was proposed.

### Four-partner interaction assays

The Protein Mimetics Library (PML-8960; Enamine), comprising 8,960 compounds, was screened at 10 µM in separate TR-FRET assays measuring LAG-3 interactions with tau, Aβ, MHC class II, and FGL1. In the FGL1 assay configuration, human Fc-tagged LAG-3 (from ACROBiosystems, Catalog# LA3-H5255) and His-tagged FGL1 (from SinoBiological, Catalog# 13484-H27B) were detected using a terbium-cryptate-labelled anti-6His monoclonal antibody donor (HTRF MAb Anti-6HIS-Tb cryptate Gold; Revvity) and an XL665-labelled anti-human IgG polyclonal antibody acceptor (HTRF PAb Anti-Human IgG-XL665; Revvity). Binding between Fc-tagged LAG-3 and His-tagged FGL1 was detected using anti-His-terbium cryptate and anti-human IgG-XL665 reagents. TR-FRET assays were performed in medium-binding Revvity OptiPlate-384 white microplate (Part #6007290, Revvity). Donor and acceptor emissions were measured on Tecan SPARK plate reader at 620 and 665 nm, respectively, following excitation at 340 nm. The TR-FRET signal was calculated as 100 × F665/F620 and normalized to vehicle-treated controls. Separate assays paired Fc-tagged LAG-3 with His-tagged tau (from ACROBiosystems, Catalog # TAU-H51H3), His-tagged Aβ42 (from ACROBiosystems, Catalog #APP-H51H6), or His-tagged MHC class II. Each interaction was detected using the anti-His– terbium cryptate donor and anti-human IgG-XL665 acceptor. Technical replicates (n=3) were averaged within each experiment. Compounds met the interaction-selectivity criterion when both disruption signals were below 50% of vehicle and both preservation signals remained within 80-120% of vehicle; prospective advancement required these criteria to be satisfied in every independent experiment.

### Orthogonal binding and interference counterscreens

Direct compound binding to LAG-3 was confirmed by SPR on a Biacore 8K. Recombinant His-tagged LAG-3 extracellular domain (10 μg/mL in PBS, pH 7.4) was immobilized on a CM5 sensor surface by amine coupling at 10 μL/min for 420 s, followed by ethanolamine blocking; the reference flow cell was mock-coupled. Compounds were injected at increasing concentrations at 30 μL/min with 120-s contact per concentration, followed by a dissociation phase. Blank injections and the reference flow cell were used for double referencing, and a DMSO calibration series was used for solvent correction. Equilibrium binding responses were plotted against compound concentration and fitted to a one-site binding model to estimate K_D_. Assays were repeated in independent experiments with technical replicates (n=3).

### Cellular tau-seeding, amyloid-β uptake, and viability

Tau-seeding activity was quantified using Tau RD P301S CFP/YFP biosensor cells (ATCC CRL-3275), which express fluorescently tagged tau repeat-domain reporters that aggregate following exposure to exogenous tau seeds. Cells were cultured in DMEM containing 10% fetal bovine serum, 1% GlutaMAX, and 1% penicillin-streptomycin at 37°C and 5% CO_2_. Cultures were confirmed free of mycoplasma. Cells were seeded at 35,000 cells per well in 96-well plates 24 h before treatment. Tau seed material, normalized to 10 µg total protein per well, was mixed with 1.5 µL Lipofectamine 2000 in Opti-MEM and incubated for 20 min at room temperature before addition to cells. Compounds were evaluated over an increasing concentration series (n=3).

After 48 h, cells were detached with TrypLE Express, washed, and analyzed by flow cytometry. CFP and YFP were excited at 405 and 488 nm, respectively; sensitized FRET emission was recorded near 525 nm following excitation at 405 nm. Cells were identified using forward- and side-scatter characteristics, followed by singlet and viability gating. CFP-only and YFP-only controls were used for compensation, and the FRET-positive gate was established using unseeded cells exposed to Lipofectamine alone. Integrated FRET density was calculated as the percentage of FRET-positive cells multiplied by their median FRET intensity. Concentration-response curves were fitted by four-parameter logistic regression for an EC_50_ estimate. Each concentration was tested in three technical wells in two independent experiments.

Cellular Aβ uptake was assessed using HiLyte Fluor 555-labelled Aβ42 (AnaSpec Inc., Catalog # AS-60480-01). Peptides were dissolved in DMSO, diluted to a total concentration of 100 µM in serum-free, phenol-red-free F-12 medium, and incubated at 4°C for 24 h to promote oligomer formation. Cryopreserved primary mouse cortical neurons isolated from embryonic day 17 C57BL/6 mice (Gibco, A15586) were used in this experiment. Cells were thawed, plated on poly-D-lysine-coated plates, and maintained according to the supplier’s instructions. Neurons were initially plated at 40,000 viable cells per well in poly-D-lysine-coated 96-well plates and maintained in 100 µL medium until DIV14. Compounds, over an eight-point concentration series, were added 1 h before exposure to 500 nM fluorescently labelled Aβ42 for 4 h at 37°C and retained throughout the exposure period. Cells were washed with PBS, and extracellular fluorescence was quenched with trypan blue before measurement of cell-associated fluorescence using a plate reader. Background-corrected fluorescence was normalized to cell abundance and expressed relative to vehicle-treated cells exposed to labelled Aβ42. Concentration-response curves were fitted by four-parameter logistic regression to estimate the concentration producing half-maximal inhibition of uptake. Viability was assessed in matched parallel wells after the same compound-exposure period. Plates were equilibrated to room temperature, and CellTiter-Glo reagent was added at a volume equal to the culture medium. Plates were shaken for 2 min and incubated for 10 min at room temperature before luminescence measurement. Background-subtracted signals were expressed as a percentage of vehicle controls. Each concentration was tested in three technical wells across two independent neuronal preparations. Cellular advancement required both tau-seeding and amyloid-β-uptake EC_50_ values below 10 µM and viability above 85%, evaluated separately in each independent experiment.

### Alzheimer’s disease donor-derived assays

Human iPSC-derived cortical neurons from three independent Alzheimer’s disease donors, obtained from FUJIFILM Cellular Dynamics (International, Madison, WI, USA), were cultured on poly-D-lysine/laminin-coated plates and matured for approximately 4 weeks. Samples were obtained from three donors with sporadic Alzheimer’s disease: a 68-year-old woman, a 74-year-old man, and a 79-year-old woman. BVX-01 was tested at 0.1, 0.3, 1, 3, and 10 µM against matched vehicle controls. Neurons were pretreated for 1 h before separate tau and Aβ42 challenges, with compound retained during exposure. Recombinant tau preformed fibrils were dispersed by sonication immediately before addition at 10 nM monomer equivalent. After 48 h, neurons were washed and harvested, and protein-normalized lysates were applied to Tau RD P301S CFP/YFP biosensor cells. Controls assessed residual extracellular seeds and compound carryover. For the Aβ42 challenge, HiLyte Fluor 555-labelled Aβ42 was diluted to 100 µM total peptide in serum-free, phenol-red-free F-12 medium, and incubated at 4°C for 24 h to promote oligomer formation. Neurons were exposed to 500 nM total Aβ42 for 4 h and analyzed immediately after washing and extracellular fluorescence quenching. Intracellular fluorescence was normalized to viable neuron number. Viability and membrane integrity were assessed in matched parallel wells using CellTiter-Glo and LDH-release assays. Each condition included three technical wells in two independent culture batches per donor. At 10 µM, advancement required both tau-seeding and Aβ42-uptake signals below 75% of matched vehicle in every donor.

### LAG-3 Binding-site mapping

Wild-type human LAG-3 extracellular domain (residues 23-450), three D1 deletion constructs (Δ52-80, Δ81-109, and Δ110-138), and seven point mutants (H56A, Y77F, R103A, V104E, R113E, Q117K, and V120D) were expressed in HEK293 cells with a C-terminal His6 tag. All constructs were verified by DNA sequencing. Proteins were purified by nickel-affinity chromatography followed by size-exclusion chromatography and exchanged into PBS, pH 7.4. Three independent expression and purification batches were generated per construct. Binding affinities for BVX-01, DN-05, and DN-07 were determined by SPR using the immobilization, referencing, and solvent-correction procedures described above. Concentration ranges were extended where necessary to characterize weaker binding to mutant constructs. Measurements were performed in technical duplicate, and affinity ratios were calculated relative to the matched wild-type preparation for each compound. Binding responses that did not support a reliable affinity estimate were reported as bounds rather than fitted values.

Functional inhibition was measured using the four-partner TR-FRET assays at increasing compound concentrations, with vehicle concentration held constant. Signals were normalized to vehicle-treated wells for the same construct. Basal binding to each partner was measured without compound and expressed relative to wild type. Construct integrity was assessed by differential scanning fluorimetry using 2 µM protein and 5× SYPRO Orange in PBS, with heating from 25 to 95°C at 1°C/min. Apparent melting temperatures were obtained from the fluorescence-transition derivative. Monomer content was assessed by analytical size-exclusion chromatography using a Superdex 200 Increase 10/300 GL column in PBS at 0.5 mL/min and calculated from integrated absorbance at 280 nm. Technical replicates were averaged within each independent protein preparation. Deletion-dependent changes in affinity and inhibition were interpreted alongside thermal stability, monomer content, and basal partner binding.

### Mouse pharmacokinetics and unbound brain exposure

Pharmacokinetics were evaluated in 10-12-week-old C57BL/6J mice after a single oral dose of BVX-01 at 10 mg/kg. BVX-01 was formulated in 0.5% methylcellulose and 0.1% Tween-80 in purified water and administered at 10 mL/kg. Three mice were terminally sampled at each of 0.5, 1, 2, 4, and 8 h. Blood was collected into EDTA-containing tubes and centrifuged at 2,000 × g for 10 min at 4°C within 30 min of collection. Mice underwent transcardial perfusion with ice-cold PBS before brain collection. Brains were weighed and homogenized in four volumes of ice-cold water. Samples were stored at −80°C until analysis.

Plasma and brain samples were extracted using acetonitrile containing an internal standard and centrifuged at 15,000 × g for 10 min. Supernatants were analyzed by LC-MS/MS using matrix-matched calibration and quality-control samples. Total brain concentrations were corrected for homogenate dilution. Fivefold-diluted brain homogenate containing 1 µM BVX-01 was dialyzed against PBS, pH 7.4, for 4 h at 37°C with shaking, in technical triplicate. Equilibration, compound stability, and recovery were assessed before calculating the unbound fraction. The measured fraction was corrected for homogenate dilution, and unbound brain concentrations were calculated as total brain concentration multiplied by the corrected unbound fraction. Peak unbound brain concentration was converted to molar units and divided by the cellular tau-seeding EC₅₀ to calculate the peak exposure-to-potency ratio.

### In vivo efficacy in PS19 and 5xFAD mice

Animal procedures were conducted under institutional IACUC protocols 2024-0010 and 2024-0006 approved by Weill Cornell Medicine. Within each model, mice were block-randomized by cage and sex to vehicle or BVX-01, with 12 animals per arm comprising six males and six females. Outcome assessors were blinded to treatment. Two independent cohorts were evaluated per model.

PS19 mice (The Jackson Laboratory, stock 008169) received BVX-01 at 30 mg/kg by once-daily oral gavage for 56 days beginning at approximately 6 months of age. Four-month-old hemizygous 5xFAD mice on a C57BL/6 background received 25 mg/kg once daily for 21 days. BVX-01 was formulated in 0.5% methylcellulose and 0.1% Tween-80 in purified water and administered at 10 mL/kg; controls received matched vehicle. Dose volumes were adjusted to body weight. Clinical condition was monitored daily and body weight at least twice weekly, with humane endpoints governed by the approved protocol.

Terminal samples were collected 2 h after the final dose. Hippocampi were collected for the PS19 tau endpoint and cortex for the 5xFAD Aβ42 endpoint. Tissue was snap-frozen and stored at −80°C, except material processed immediately for thermal-shift measurements.

#### Hippocampal tau aggregation

Hippocampi were homogenized in ten volumes of ice-cold 20 mM HEPES, pH 7.4, 150 mM NaCl, and 1% Triton X-100 containing protease and phosphatase inhibitors. Homogenates were centrifuged at 20,000 × g for 20 min at 4°C. Supernatant protein concentrations were measured by a detergent-compatible assay. Tau aggregation-associated signal was quantified using the HTRF Tau Aggregation Kit (Revvity), with equal protein inputs and sample dilutions established to avoid saturation and the hook effect. Each 20-µL reaction contained 10 µL sample and 5 µL each of the kit’s anti-tau terbium-cryptate and d2 working reagents. After 2 h at room temperature, emissions were measured at 620 and 665 nm. Background-corrected acceptor-to-donor ratios were averaged across technical duplicates and expressed relative to the matched vehicle group.

#### Cortical Aβ42

Cortex was homogenized in ten volumes of ice-cold TBS containing protease inhibitors and centrifuged at 100,000 × g for 60 min at 4°C. The TBS-insoluble pellet was resuspended in ten volumes, relative to starting tissue mass, of 5 M guanidine hydrochloride in 50 mM Tris-HCl, pH 8.0. Samples were mixed for 3 h at room temperature and centrifuged at 20,000 × g for 20 min. Supernatants were diluted into the validated working range of the Human Aβ42 ELISA Kit (Invitrogen, KHB3441). Concentrations were interpolated from the calibration curve, corrected for extraction and assay dilution, and expressed as nanograms per milligram of starting wet cortical tissue. Technical duplicates were averaged to obtain one value per animal. Cortical TBS-insoluble, guanidine-extractable Aβ42 constituted the sole amyloid efficacy endpoint.

Measurements were analyzed separately within each model and cohort using a two-sided Mann-Whitney U test with the asymptotic approximation at α = 0.05. Advancement required the pathology endpoint to be no more than 70% of vehicle with P < 0.05 in both models. Body-weight retention was calculated relative to pretreatment baseline.

#### Brain LAG-3 thermal-shift assay

Fresh brain tissue was homogenized in 20 mM HEPES, pH 7.4, 150 mM NaCl, and 0.2% NP-40 substitute containing protease and phosphatase inhibitors. A preliminary temperature-response experiment established the LAG-3 thermal challenge. Matched aliquots were retained unheated or heated for 3 min at the selected temperature, cooled for 3 min at room temperature, and centrifuged at 20,000 × g for 20 min at 4°C. Supernatant LAG-3 was quantified by sandwich immunoassay within its validated working range. Heated signals were normalized to matched unheated inputs and expressed relative to vehicle controls. These measurements were reported as thermal-shift responses.

### Chemical synthesis and characterization

De novo compounds were synthesized by amide coupling of the carboxylic acid and amine building blocks listed in table S2. The acid (1.0 equiv), amine (1.1 equiv), HATU (1.2 equiv), and DIPEA (3.0 equiv) were combined in anhydrous DMF at 0.1 M and stirred at room temperature for 4-16 h. Reaction progress was monitored by LC-MS. Products were purified by flash column chromatography using a hexane-ethyl acetate gradient. Purity was determined by analytical reversed-phase HPLC with UV detection and confirmed to meet ≥95% before testing. All synthesized compounds were characterized using high-resolution mass spectrometry using Waters Acquity Xevo G3. HRMS data for each compound are given in figs. S9 to S17

### Cross-target biochemical assays for PrP^c^ and LRP1

The PrPᶜ panel measured interactions with Aβ42 oligomers and tau oligomers, designated for disruption, and with STI1 and laminin, designated for preservation. The LRP1 panel measured interactions with tau and APP751-770, designated for disruption, and with APOE3 and activated α₂- macroglobulin, designated for preservation. Reagents included full-length STI1 (OriGene), mouse laminin-111 (Sigma-Aldrich), human LRP1 cluster II–Fc (R&D Systems, 2368-L2), human APOE3 (R&D Systems, 4144-AE), and human plasma α₂-macroglobulin (Sigma-Aldrich, M6159), subjected to activation before testing. Tau and Aβ42 preparations matched those used in the corresponding biochemical assays.

Interactions were measured separately by TR-FRET using biotinylated receptor and streptavidin– terbium cryptate (Revvity, 610SATLF), paired with an acceptor-labelled partner or partner-specific detection reagent. Reactions contained 20 µL PBS, pH 7.4, with 0.1% BSA and 0.01% Tween-20; LRP1 assays additionally contained 1 mM CaCl_2_. Compounds were tested at 10 µM and 0.1% DMSO, with a 30-min receptor preincubation followed by a 2-h incubation with partner at room temperature. Emissions at 620 and 665 nm were used to calculate background-corrected ratios normalized to vehicle controls. Agent-selected and random-control compounds were tested together in technical triplicate across two independent experiments. Hits required both disruption signals below 50% of vehicle and both preservation signals within 80-120% in each experiment.

### Statistics and reporting

Prospective arm comparisons on the ported targets use Fisher’s exact test with the direction stated in advance. The chemist comparison is reported descriptively: no comparison statistic, pooling scheme or success threshold was fixed before those data were seen, so counts and Clopper-Pearson intervals are given and no hypothesis test is reported. This differs from the ported-target comparisons, where the arms and the agent-versus-random comparison were prespecified and only the pooling across targets was post hoc. Counts against a library base rate use the exact binomial test; in vivo comparisons use the two-sided Mann-Whitney U test with the asymptotic approximation used by the runner. Binomial confidence intervals are Wilson intervals. Cross-validation holds out whole Bemis-Murcko scaffolds in five folds.

## Supplementary Materials

fig. S1. SPR binding isotherms for compounds advanced to orthogonal confirmation.

fig. S2. Concentration-response relationships for the compounds advancing to cellular assays.

fig. S3. The nine de novo compounds.

fig. S4. SPR binding isotherms for the de novo compounds carried to orthogonal confirmation.

fig. S5. Dissociation constants per construct for each compound.

fig. S6. Construct validation for the binding-site mapping.

fig. S7. Sequential closed-loop campaigns on PrPᶜ and LRP1.

fig. S8. Near-neighbour control for the ported campaigns.

figs. S9 to S17. HRMS data for compounds DN-01 to DN-09.

table S1. Prospective selection commitments.

table S2. The de novo series and its building blocks.

Supplementary Methods

## Author contributions

The manuscript was written through contributions of all authors. All authors have given approval to the final version of the manuscript. conceptualization: M.T.G. and S.A.A.-R. methodology: S.A.A.-R. and M.T.G. investigation: S.A.A.-R. and M.T.G. formal analysis: S.A.A.-R. and M.T.G. visualization: S.A.A.-R. and M.T.G. supervision: M.T.G. writing-original draft: M.T.G. and S.A.A.-R. writing-review and editing: M.T.G. and S.A.A.-R. data curation: M.T.G. validation: M.T.G. project administration: M.T.G.

## REFERENCES

1. A. Zhavoronkov, Y. A. Ivanenkov, A. Aliper, M. S. Veselov, V. A. Aladinskiy, A. V. Aladinskaya, V. A. Terentiev, D. A. Polykovskiy, M. D. Kuznetsov, A. Asadulaev, Y. Volkov, A. Zholus, R. R. Shayakhmetov, A. Zhebrak, L. I. Minaeva, B. A. Zagribelnyy, L. H. Lee, R. Soll, D. Madge, L. Xing, T. Guo, A. Aspuru-Guzik, Deep learning enables rapid identification of potent DDR1 kinase inhibitors. Nat. Biotechnol. 37, 1038–1040 (2019). doi:10.1038/s41587-019-0224-x

2. J. M. Stokes, K. Yang, K. Swanson, W. Jin, A. Cubillos-Ruiz, N. M. Donghia, C. R. MacNair, S. French, L. A. Carfrae, Z. Bloom-Ackermann, V. M. Tran, A. Chiappino-Pepe, A. H. Badran, I. W. Andrews, E. J. Chory, G. M. Church, E. D. Brown, T. S. Jaakkola, R. Barzilay, J. J. Collins, A deep learning approach to antibiotic discovery. Cell 180, 688–702 (2020). doi:10.1016/j.cell.2020.01.021

3. F. Wong, E. J. Zheng, J. A. Valeri, N. M. Donghia, M. N. Anahtar, S. Omori, A. Li, A. Cubillos-Ruiz, A. Krishnan, W. Jin, A. L. Manson, J. Friedrichs, R. Helbig, B. Hajian, D. K. Fiejtek, F. F. Wagner, H. H. Soutter, A. M. Earl, J. M. Stokes, L. D. Renner, J. J. Collins, Discovery of a structural class of antibiotics with explainable deep learning. Nature 626, 177–185 (2024). doi:10.1038/s41586-023-06887-8

4. J. Jumper, R. Evans, A. Pritzel, T. Green, M. Figurnov, O. Ronneberger, K. Tunyasuvunakool, R. Bates, A. Ží dek, A. Potapenko, A. Bridgland, C. Meyer, S. A. A. Kohl, A. J. Ballard, A. Cowie, B. Romera-Paredes, S. Nikolov, R. Jain, J. Adler, T. Back, S. Petersen, D. Reiman, E. Clancy, M. Zielinski, M. Steinegger, M. Pacholska, T. Berghammer, S. Bodenstein, D. Silver, O. Vinyals, A. W. Senior, K. Kavukcuoglu, P. Kohli, D. Hassabis, Highly accurate protein structure prediction with AlphaFold. Nature 596, 583–589 (2021). doi:10.1038/s41586-021-03819-2

5. F. Ren, A. Aliper, J. Chen, H. Zhao, S. Rao, C. Kuppe, I. V. Ozerov, M. Zhang, K. Witte, C. Kruse, V. Aladinskiy, Y. Ivanenkov, D. Polykovskiy, Y. Fu, E. Babin, J. Qiao, X. Liang, Z. Mou, H. Wang, F. W. Pun, P. Torres-Ayuso, M. Veviorskiy, D. Song, S. Liu, B. Zhang, V. Naumov, X. Ding, A. Kukharenko, E. Izumchenko, A. Zhavoronkov, A small-molecule TNIK inhibitor targets fibrosis in preclinical and clinical models. Nat. Biotechnol. 43, 63–75 (2025). doi:10.1038/s41587-024-02143-0

6. A. E. Ghareeb, B. Chang, L. Mitchener, A. Yiu, C. Szostkiewicz, D. Shved, G. Gyimesi, J. Laurent, S. Wright, M. Razzak, A. White, S. Finnemann, M. Hinks, S. Rodriques, A multi-agent system for automating scientific discovery. Nature, 655, 497–505 (2026). 10.1038/s41586-026-10652-y

7. S. Kapoor, A. Narayanan, Leakage and the reproducibility crisis in machine-learning-based science. Patterns 4, 100804 (2023). doi:10.1016/j.patter.2023.100804

8. R. P. Sheridan, Time-split cross-validation as a method for estimating the goodness of prospective prediction. J. Chem. Inf. Model. 53, 783–790 (2013). doi:10.1021/ci400084k

9. I. Wallach, A. Heifets, Most ligand-based classification benchmarks reward memorization rather than generalization. J. Chem. Inf. Model. 58, 916–932 (2018). doi:10.1021/acs.jcim.7b00403

10. N. Brown, M. Fiscato, M. H. S. Segler, A. C. Vaucher, GuacaMol: Benchmarking models for de novo molecular design. J. Chem. Inf. Model. 59, 1096–1108 (2019). doi:10.1021/acs.jcim.8b00839

11. J. H. Jensen, A graph-based genetic algorithm and generative model/Monte Carlo tree search for the exploration of chemical space. Chem. Sci. 10, 3567–3572 (2019). doi:10.1039/C8SC05372C

12. A. Bender, I. Cortes-Ciriano, Artificial intelligence in drug discovery: What is realistic, what are illusions? Part 1: Ways to make an impact, and why we are not there yet. Drug Discov. Today 26, 511–524 (2021). doi:10.1016/j.drudis.2020.12.009

13. J. Jiménez-Luna, F. Grisoni, G. Schneider, Drug discovery with explainable artificial intelligence. Nat. Mach. Intell. 2, 573–584 (2020). doi:10.1038/s42256-020-00236-4

14. D. Stumpfe, J. Bajorath, Exploring activity cliffs in medicinal chemistry. J. Med. Chem. 55, 2932–2942 (2012). doi:10.1021/jm201706b

15. R. Muruganantham, S. K. Mohanty, A. Das, S. N. Patil, AI-augmented medicinal chemist: Skillsets for resolving uncertainty in the AI-assisted drug discovery era. J. Med. Chem. In Press (2026). doi:10.1021/acs.jmedchem.6c03403

16. E. Granqvist, R. Mercado, S. Genheden, Designing an LLM-based evaluation system for agentic AI in drug discovery through human alignment. Artif. Intell. Life Sci. 10, 100187 (2026). doi:10.1016/j.ailsci.2026.100187

17. B. A. Nosek, C. R. Ebersole, A. C. DeHaven, D. T. Mellor, The preregistration revolution. Proc. Natl. Acad. Sci. U.S.A. 115, 2600–2606 (2018). doi:10.1073/pnas.1708274114

18. A.-W. Chan, J. M. Tetzlaff, D. G. Altman, A. Laupacis, P. C. Gøtzsche, K. Krleža-Jerić, A. Hróbjartsson, H. Mann, K. Dickersin, J. A. Berlin, C. J. Doré, W. R. Parulekar, W. S. M. Summerskill, T. Groves, K. F. Schulz, H. C. Sox, F. W. Rockhold, D. Rennie, D. Moher, SPIRIT 2013 statement: Defining standard protocol items for clinical trials. Ann. Intern. Med. 158, 200–207 (2013). doi:10.7326/0003-4819-158-3-201302050-00583

19. B. Huard, P. Prigent, M. Tournier, D. Bruniquel, F. Triebel, CD4/major histocompatibility complex class II interaction analyzed with CD4- and lymphocyte activation gene-3 (LAG-3)-Ig fusion proteins. Eur. J. Immunol. 25, 2718–2721 (1995). doi:10.1002/eji.1830250949

20. T. Maruhashi, D. Sugiura, I. Okazaki, T. Okazaki, LAG-3: From molecular functions to clinical applications. J. Immunother. Cancer 8, e001014 (2020). doi:10.1136/jitc-2020-001014

21. J. Wang, M. F. Sanmamed, I. Datar, T. T. Su, L. Ji, J. Sun, L. Chen, Y. Chen, G. Zhu, W. Yin, L. Zheng, T. Zhou, T. Badri, S. Yao, S. Zhu, A. Boto, M. Sznol, I. Melero, D. A. A. Vignali, K. Schalper, L. Chen, Fibrinogen-like protein 1 is a major immune inhibitory ligand of LAG-3. Cell 176, 334–347 (2019). doi:10.1016/j.cell.2018.11.010

22. X. Mao, M. T. Ou, S. S. Karuppagounder, T.-I. Kam, X. Yin, Y. Xiong, P. Ge, G. E. Umanah, S. Brahmachari, J.-H. Shin, H. C. Kang, J. Zhang, J. Xu, R. Chen, H. Park, S. A. Andrabi, S. U. Kang, R. A. Gonçalves, Y. Liang, S. Zhang, C. Qi, S. Lam, J. A. Keiler, J. Tyson, D. Kim, N. Panicker, S. P. Yun, C. J. Workman, D. A. A. Vignali, V. L. Dawson, H. S. Ko, T. M. Dawson, Pathological α-synuclein transmission initiated by binding lymphocyte-activation gene 3. Science 353, aah3374 (2016). doi:10.1126/science.aah3374

23. M. Emmenegger, E. De Cecco, M. Hruska-Plochan, T. Eninger, M. Schneider, M. Barth, E. Tantardini, P. de Rossi, M. Bacioglu, R. Langston, A. Kaganovich, N. Bengoa-Vergniory, A. Gonzalez-Guerra, M. Avar, D. Heinzer, R. Reimann, L. Häsler, T. Herling, N. Matharu, N. Landeck, K. Luk, R. Melki, P. Kahle, S. Hornemann, T. Knowles, M. Cookson, M. Polymenidou, M. Jucker, A. Aguzzi, LAG3 is not expressed in human and murine neurons and does not modulate α-synucleinopathies. EMBO Mol. Med. 13, e14745 (2021). doi:10.15252/emmm.202114745

24. C. Chen, R. Kumbhar, H. Wang, X. Yang, K. Gadhave, C. Rastegar, Y. Kimura, A. Behensky, S. Kotha, G. Kuo, S. Katakam, D. Jeong, L. Wang, A. Wang, R. Chen, S. Zhang, L. Jin, C. Workman, D. Vignali, O. Pletinkova, H. Jia, W. Peng, D. Nauen, P. Wong, J. Redding-Ochoa, J. Troncoso, M. Ying, V. Dawson, T. Dawson, X. Mao, Lymphocyte-activation gene 3 facilitates pathological tau neuron-to-neuron transmission. Adv. Sci. 11, 2303775 (2024). doi:10.1002/advs.202303775

25. J. M. Long, D. M. Holtzman, Alzheimer disease: An update on pathobiology and treatment strategies. Cell 179, 312–339 (2019). doi:10.1016/j.cell.2019.09.001

26. E. E. Congdon, C. Ji, A. M. Tetlow, Y. Jiang, E. M. Sigurdsson, Tau-targeting therapies for Alzheimer disease: Current status and future directions. Nat. Rev. Neurol. 19, 715–736 (2023). doi:10.1038/s41582-023-00883-2

27. Y. Yoshiyama, M. Higuchi, B. Zhang, S.-M. Huang, N. Iwata, T. C. Saido, J. Maeda, T. Suhara, J. Q. Trojanowski, V. M.-Y. Lee, Synapse loss and microglial activation precede tangles in a P301S tauopathy mouse model. Neuron 53, 337–351 (2007). doi:10.1016/j.neuron.2007.01.010

28. H. Oakley, S. L. Cole, S. Logan, E. Maus, P. Shao, J. Craft, A. Guillozet-Bongaarts, M. Ohno, J. Disterhoft, L. Van Eldik, R. Berry, R. Vassar, Intraneuronal β-amyloid aggregates, neurodegeneration, and neuron loss in transgenic mice with five familial Alzheimer’s disease mutations: potential factors in amyloid plaque formation. J. Neurosci. 26, 10129–10140 (2006). doi:10.1523/JNEUROSCI.1202-06.2006

29. J. Laurén, D. A. Gimbel, H. B. Nygaard, J. W. Gilbert, S. M. Strittmatter, Cellular prion protein mediates impairment of synaptic plasticity by amyloid-β oligomers. Nature 457, 1128–1132 (2009). doi:10.1038/nature07761

30. J. N. Rauch, G. Luna, E. Guzman, M. Audouard, C. Challis, Y. E. Sibih, C. Leshuk, I. Hernandez, S. Wegmann, B. T. Hyman, V. Gradinaru, M. Kampmann, K. S. Kosik, LRP1 is a master regulator of tau uptake and spread. Nature 580, 381–385 (2020). doi:10.1038/s41586-020-2156-5

31. S. A. Abdel-Rahman, M. T. Gabr, HTS-Oracle v2: Prospective Ai-Guided Discovery and Experimental Validation of Small Molecule Modulators Across Multiple Targets. J. Chem. Inf. Model. 66, 8998–9008 (2026). 10.1021/acs.jcim.6c01522

32. S. A. Abdel-Rahman, M. T. Gabr, HTS-Oracle X: AI-Guided Prospective Discovery of Small Molecule Immune Checkpoint Binders. ACS Med Chem Lett. 17, 1888–1894 (2026). doi:10.1021/acsmedchemlett.6c00289

33. S. A. Abdel-Rahman, M. T. Gabr, OracleScreen-LILRB4 (HTS-Oracle v3): machine learning-guided discovery of myeloid immune checkpoint binders validated in patient-derived cells. Bioorg. Med. Chem. Lett. 140, 130717 (2026). doi:10.1016/j.bmcl.2026.130717

34. G. W. Bemis, M. A. Murcko, The properties of known drugs. 1. Molecular frameworks. J. Med. Chem. 39, 2887–2893 (1996). doi:10.1021/jm9602928

35. G. A. Landrum, S. Riniker, Combining IC50 or Ki values from different sources is a source of significant noise. J. Chem. Inf. Model. 64, 1560–1567 (2024). doi:10.1021/acs.jcim.4c00049

36. D. Rogers, M. Hahn, Extended-connectivity fingerprints. J. Chem. Inf. Model. 50, 742–754 (2010). doi:10.1021/ci100050t

37. G. Landrum, RDKit: Open-source cheminformatics (2024); www.rdkit.org.

38. F. Pedregosa, G. Varoquaux, A. Gramfort, V. Michel, B. Thirion, O. Grisel, M. Blondel, P. Prettenhofer, R. Weiss, V. Dubourg, J. Vanderplas, A. Passos, D. Cournapeau, M. Brucher, M. Perrot, E. Duchesnay, Scikit-learn: Machine learning in Python. J. Mach. Learn. Res. 12, 2825–2830 (2011).

