## Supporting Information for "An AI agent guides a pipeline for selective protein interaction inhibitors from discovery across targets to preclinical evaluation"

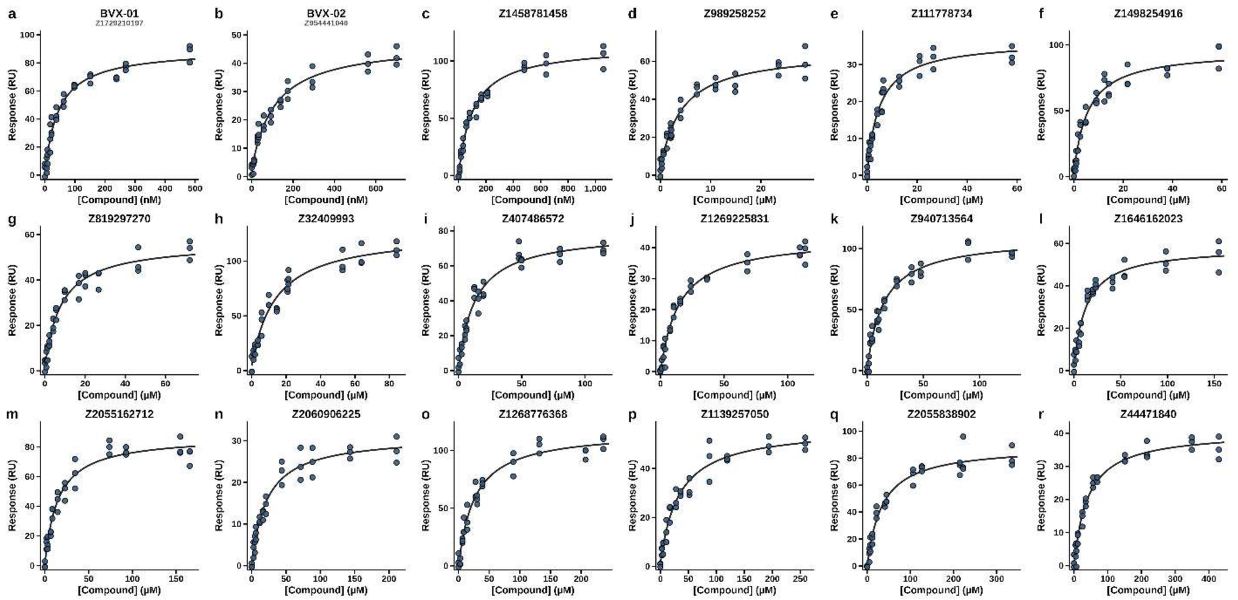


**Fig. S1. Surface plasmon resonance binding isotherms for all compounds advanced to orthogonal confirmation.** Equilibrium response against compound concentration for the 18 compounds that passed the primary biochemical gate, ordered by affinity. **a**, BVX-01 (Z1729210197). **b**, BVX-02 (Z954441040). **c-r**, the remaining compounds by catalogue identifier. Every replicate is plotted; curves are fits to a one-site binding model from which dissociation constants were derived. Dissociation constants were measured in two independent experiments of three replicates each.

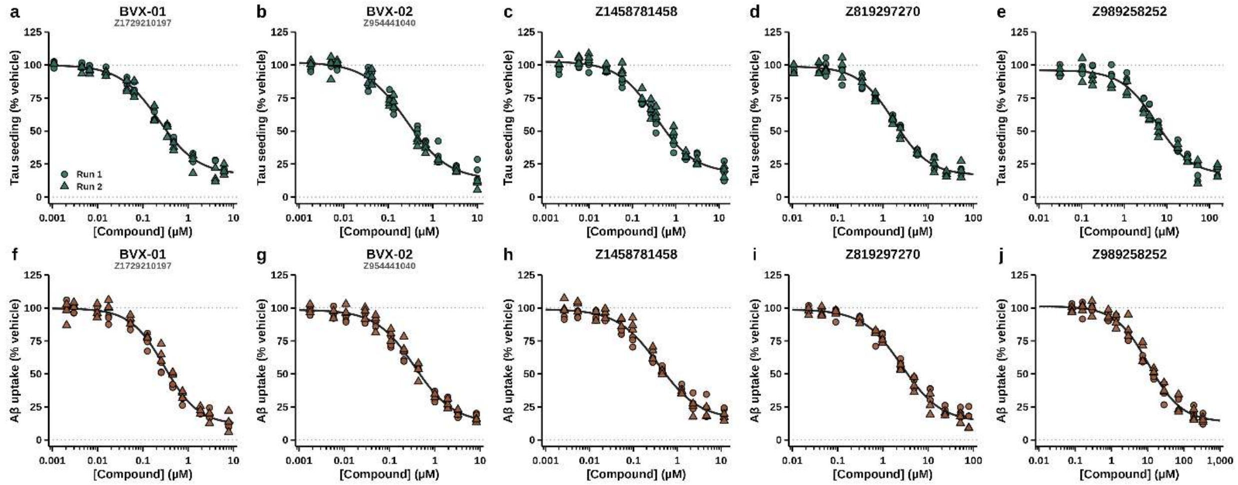


**Fig. S2. Concentration-response relationships for the five compounds advancing to cellular assays. a-e**, Tau seeding and **f-j**, Aβ uptake, each expressed as a percentage of vehicle, for BVX-01 (Z1729210197), BVX-02 (Z954441040), Z1458781458, Z819297270 and Z989258252. Circles and triangles distinguish two independent runs and every replicate is plotted; curves are four-parameter logistic fits to the data, from which the $EC_{50}$ values were derived. Dotted lines mark the vehicle control and complete inhibition.

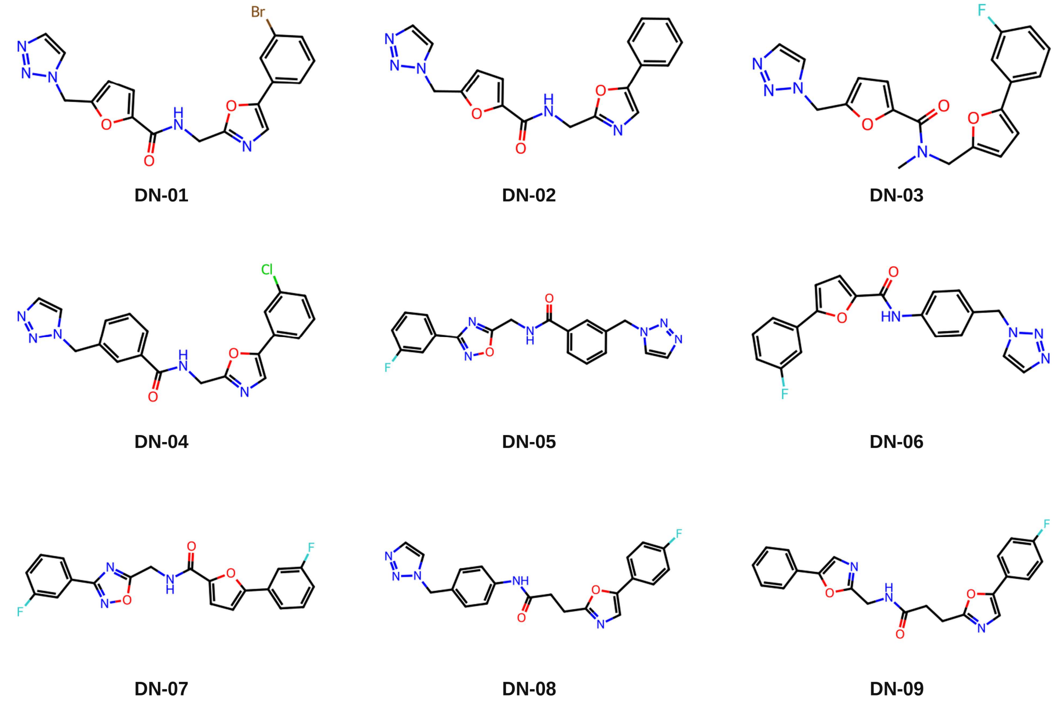


**Fig. S3. The nine de novo compounds.** Each is assembled in a single amide coupling from two commercially available building blocks; supplier codes are given in Supplementary Table 2.

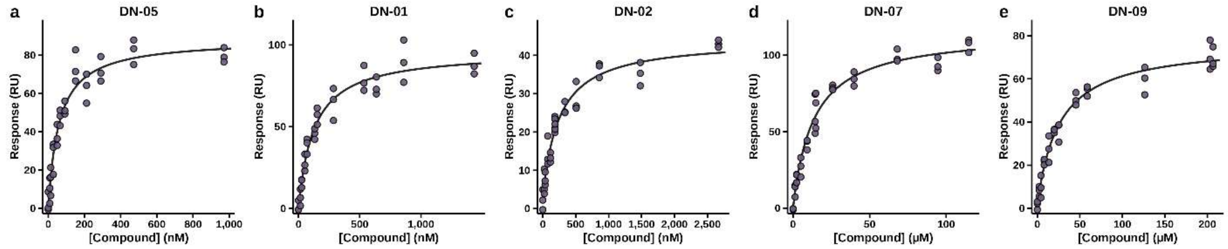


**Fig. S4. Surface plasmon resonance binding isotherms for the de novo compounds carried to orthogonal confirmation.** Equilibrium response against compound concentration, ordered by affinity. **a**, DN-05. **b**, DN-01. **c**, DN-02. **d**, DN-07. **e**, DN-09. Every replicate is plotted; curves are fits to a one-site binding model.

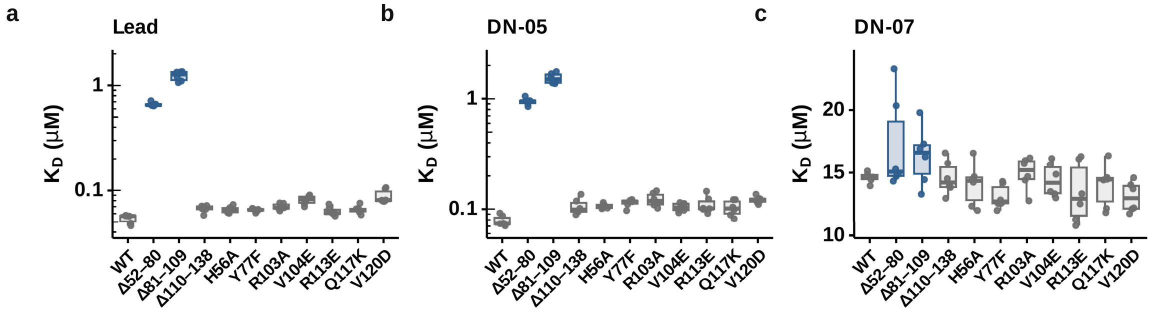


**Fig. S5. Dissociation constants per construct, underlying the summary in Fig. 4. a,** Campaign lead (BVX-01). **b,** DN-05. **c,** The triazole-free DN-07. Each point is one independent protein preparation; boxes show the median and interquartile range with whiskers to 1.5× the interquartile range, and filled boxes mark the two deletions spanning residues 52-109.

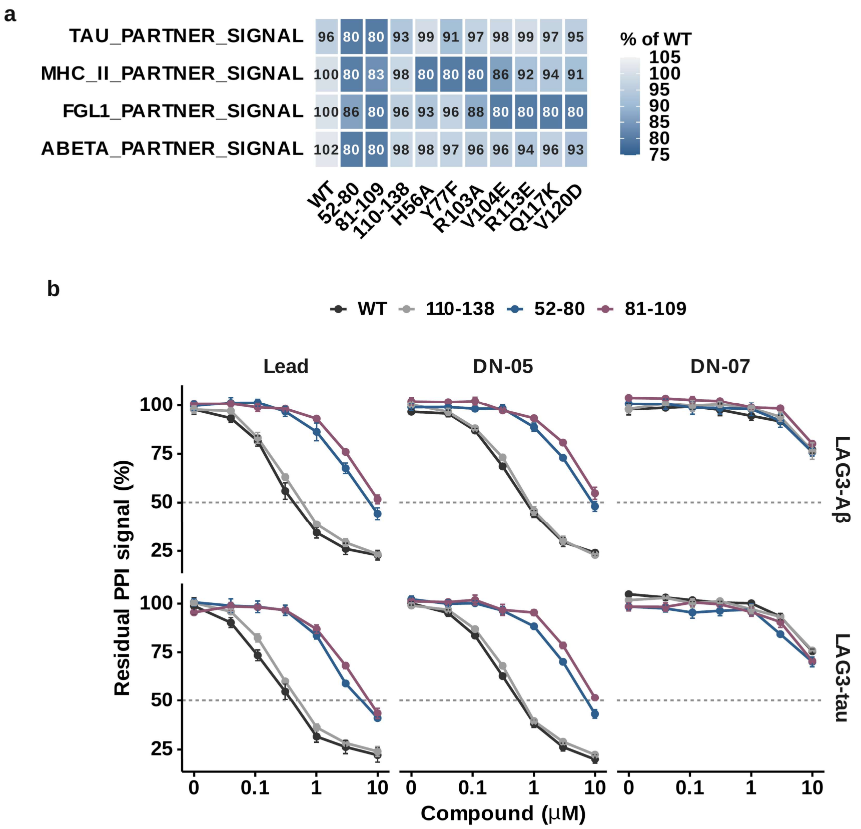


**Fig. S6. Construct validation for the binding-site mapping. a,** Basal binding of all four LAG-3 partners to every construct, expressed as a percentage of wild type and measured without compound. Each point mutant selectively reduces binding of its intended partner, confirming assay sensitivity. **b,** Dose-response for residual LAG3-tau and LAG3-Aβ signal across all three compounds (BVX-01 Lead, DN-05, and DN-07) and the deletion panel; points are means of three independent protein preparations, and error bars show s.e.m.

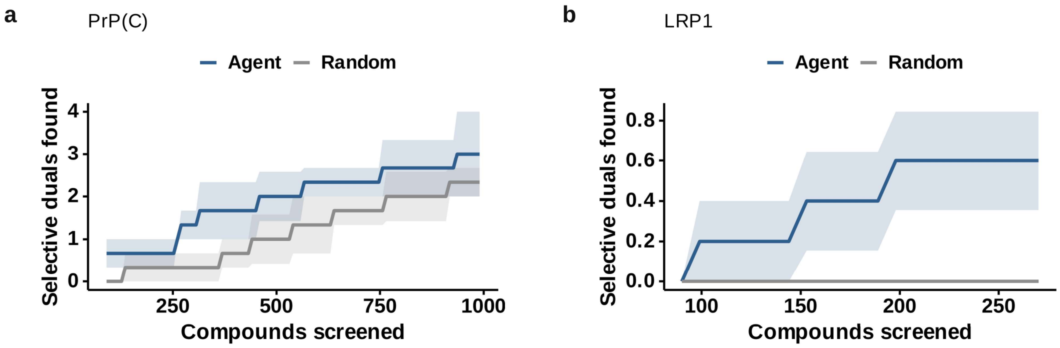


**Fig. S7. Sequential closed-loop campaigns on the ported targets. a,** PrP$^{c}$. **b,** LRP1. Cold start of 90 randomly chosen compounds followed by batches of 9 selected by the agent with retraining on every returned result, against a random arm drawn from the same seed set. Lines are means of independent seeds; shaded bands show s.e.m. Neither target shows separation from random selection at this screening depth, which preceded and contrasted with the prospective outcomes in Fig. 5.

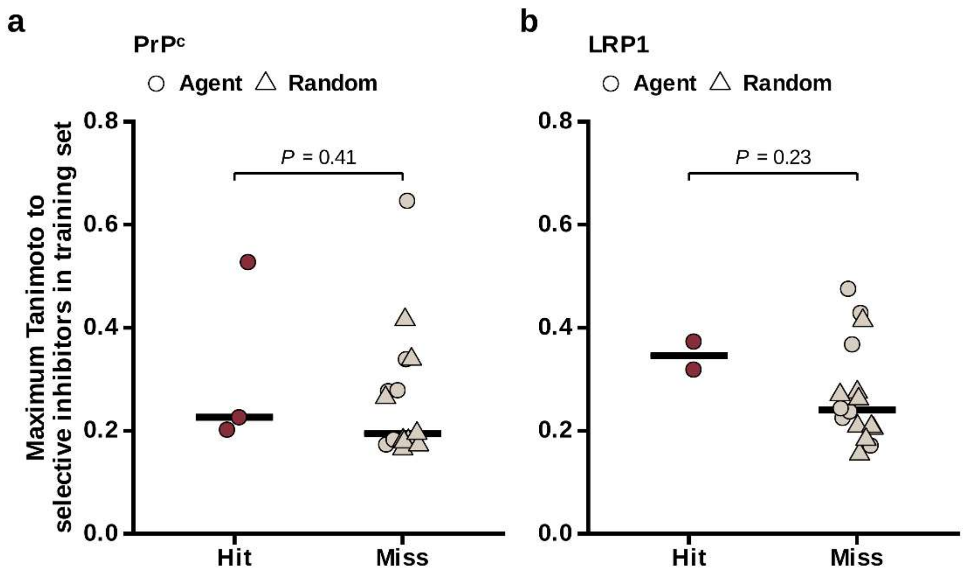


**Fig. S8.** Near-neighbor control for the ported campaigns. Maximum Tanimoto similarity (Morgan, radius 2, 2048 bits) from each prospectively screened compound to any selective inhibitor in the training set, separated into hits and misses for each target. Hits were not detectably more similar to training-set selective inhibitors than misses, although the small number of hits limits this comparison.

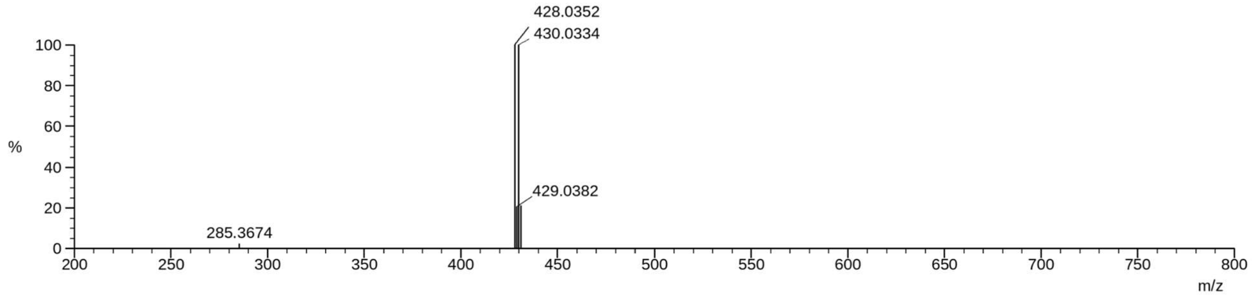


**Fig. S9.** HRMS data for compound DN-01.

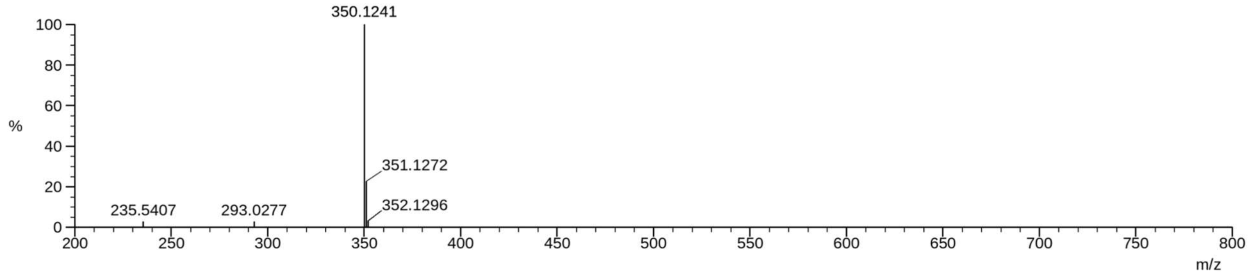


**Fig. S10.** HRMS data for compound DN-02.

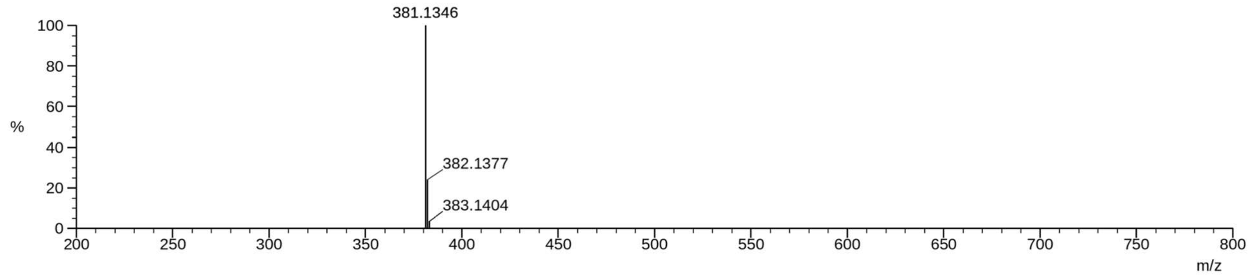


**Fig. S11.** HRMS data for compound DN-03.

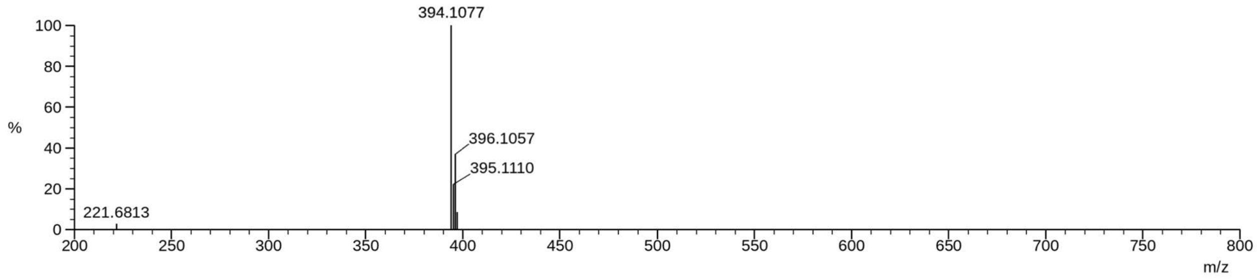


**Fig. S12.** HRMS data for compound DN-04.

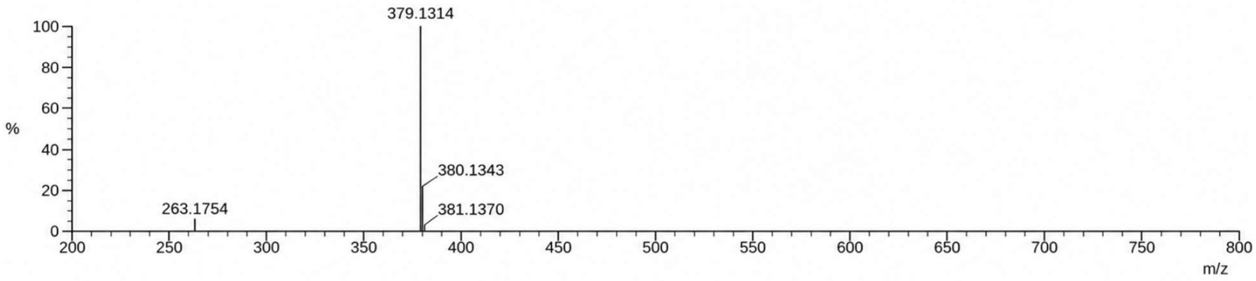


**Fig. S13.** HRMS data for compound DN-05.

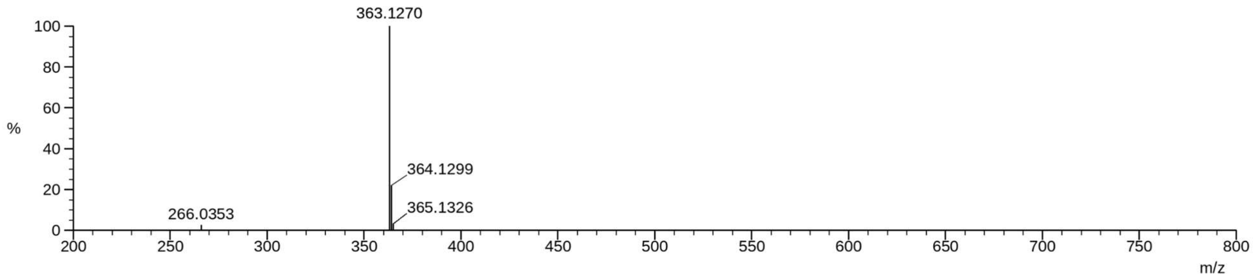


**Fig. S14.** HRMS data for compound DN-06.

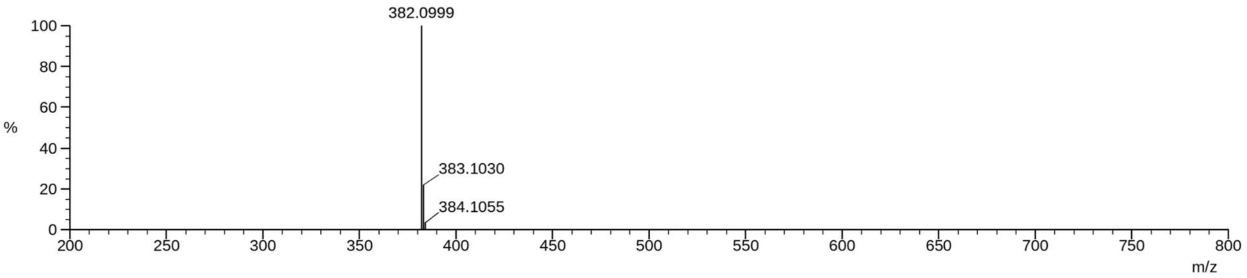


**Fig. S15.** HRMS data for compound DN-07.

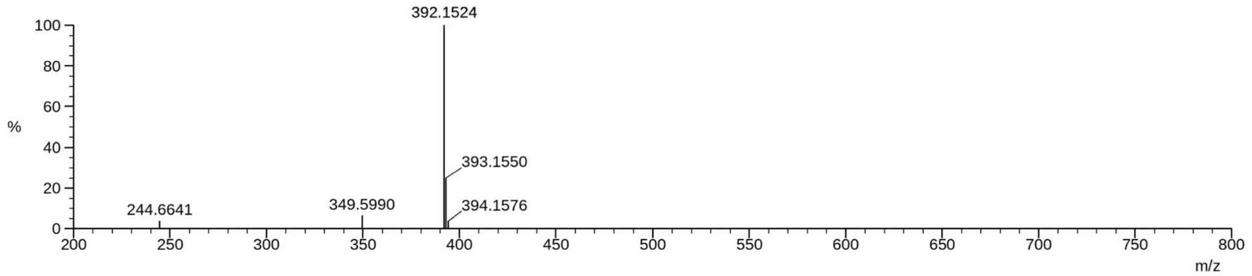


**Fig. S16.** HRMS data for compound DN-08.

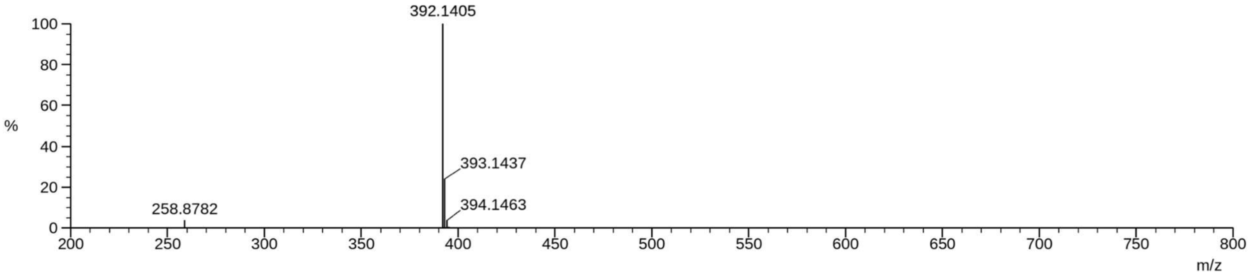


**Fig. S17.** HRMS data for compound DN-09.

**Supplementary Tables**

**Table S1. Prospective selection commitments.**

The file path in the table identifies the repository record that stores a commitment. The two chemist-arm digests were created after screening and establish the selected compound lists. To recompute a payload digest, run python3 scripts/verify_commitment_payload.py <payload.json>. Use the ID files for the agent arm and de novo rows, and the commitment JSON files for the rows whose compound_ids are included. Arm and batch digests are computed over sorted compound identifiers joined by commas; the de novo digest is computed over sorted canonical SMILES joined by commas.

| Commitment / payload hashed | Repository record (stores commitment) | Commitment SHA-256 | Timing |
|---|---|---|---|
| Sorted compound IDs, comma-joined | results/comparator/arm_agent_commitment_v3.json; payload .arm_agent_SEALED_v3.json | d04f96ef881333abefba034af74e2adf5951638ebc2129d2bff87388fdf12185 | before screening |
| Sorted compound IDs, comma-joined | results/comparator/arm_random_v3.json | 0756329c1ff0573fb7cdf22e20019d2bc7006ca5c91cef8142ffa5af63d104d7 | before screening |
| Sorted compound IDs, comma-joined | results/comparator/arm_chemist_A.json | fdac71e4a65af1a7e6ffa645219dfd8c481e40158a9d7d5178282263342c4373 | after screening |
| Sorted compound IDs, comma-joined | results/comparator/arm_chemist_B.json | 69b60401a1bf4a80b5399bfa0cce282336b65e463087143e2d27b0609b13cd75 | after screening |
| Sorted canonical SMILES, comma-joined | results/dn_series/commitment.json; payload .dn_SEALED.json | 4e2e4458706dd7d396adb27de650305059867bec5b3d61d98661f115ea426eaf | before screening |
| Sorted compound IDs, comma-joined | results/prpc_campaign/batch_01_commitment.json | 5dec781b41b8242c9d2c4b834d727406bbe89d0e6919f87291156cd13cba3f9f | before screening |
| Sorted compound IDs, comma-joined | results/lrp1_campaign/batch_01_commitment.json | 3a59e00c80e696849f1f3429df0feb2441d125f03a68ed329245910441d1200d | before screening |
| Sorted compound IDs, comma-joined | results/prpc_campaign/control_random_commitment.json | c63e3f4c253a58b6d33500ac1ba95400db17f2ee16f4e836adc4256d23b9c746 | before screening |
| Sorted compound IDs, comma-joined | results/lrp1_campaign/control_random_commitment.json | 015d8c5350f55a6328e201f6fcf8a6d8484fe25387352a50bbb0ef511eb948d0 | before screening |

**Table S2. The de novo series.**

Canonical SMILES as sealed, with the Enamine catalogue identifiers of the acid and amine building blocks from which each compound is assembled in one amide coupling.

| Compound_id | Canonical_SMILES | Acid_block_id | Amine_block_id |
|---|---|---|---|
| DN-01 | O=C(NCc1ncc(-c2cccc(Br)c2)o1)c1ccc(Cn2ccnn2)o1 | EN300-706128 | EN300-7455352 |
| DN-02 | O=C(NCc1ncc(-c2ccccc2)o1)c1ccc(Cn2ccnn2)o1 | EN300-706128 | EN300-69271 |
| DN-03 | CN(Cc1ccc(-c2cccc(F)c2)o1)C(=O)c1ccc(Cn2ccnn2)o1 | EN300-706128 | EN300-24525686 |
| DN-04 | O=C(NCc1ncc(-c2cccc(Cl)c2)o1)c1cccc(Cn2ccnn2)c1 | EN300-307078 | EN300-52423165 |
| DN-05 | O=C(NCc1nc(-c2cccc(F)c2)no1)c1cccc(Cn2ccnn2)c1 | EN300-307078 | EN300-188001 |
| DN-06 | O=C(Nc1ccc(Cn2ccnn2)cc1)c1ccc(-c2cccc(F)c2)o1 | EN300-1131868 | EN300-3007559 |
| DN-07 | O=C(NCc1nc(-c2cccc(F)c2)no1)c1ccc(-c2cccc(F)c2)o1 | EN300-1131868 | EN300-188001 |
| DN-08 | O=C(CCc1ncc(-c2ccc(F)cc2)o1)Nc1ccc(Cn2ccnn2)cc1 | EN300-26340 | EN300-3007559 |
| DN-09 | O=C(CCc1ncc(-c2ccc(F)cc2)o1)NCc1ncc(-c2ccccc2)o1 | EN300-26340 | EN300-69271 |

### Supplementary Methods

**Molecular representation and endpoint models**:

Compounds were featurised as a 1024-bit Morgan count fingerprint of radius 2 concatenated with ten RDKit descriptors: molecular weight, Crippen logP, topological polar surface area, hydrogen-bond acceptor count, hydrogen-bond donor count, rotatable-bond count, ring count, aromatic ring count, fraction of $sp^3$ carbons, and heavy-atom count (1,034 features in total). Unparsable structures yield a zero row and are excluded from selection. One RandomForestClassifier was fitted per endpoint with n_estimators = 400, min_samples_leaf = 4, max_features = "sqrt", class_weight = "balanced" and random_state = 901. Where an endpoint had a single class in the training labels, the model returned the base rate with zero uncertainty. Features are versioned as morgan1024r2+desc10/v1; a model saved under one feature version will not load under another. Similarity, used for the applicability domain, the downstream-failure exclusion and the near-neighbor control in fig. S8, was computed from 2048-bit binary Morgan fingerprints of radius 2.
